# Brr2p-mediated unwinding of U4/U6 is promoted by a mutually exclusive intra-molecular stem loop in U4 and involves destabilization of the 5’ stem-loop of U4

**DOI:** 10.1101/2025.08.01.667943

**Authors:** Klaus H Nielsen, Amartya Das, Jonathan P Staley

**Affiliations:** Department of Molecular Genetics and Cell Biology, University of Chicago, Chicago, IL 60637, USA; Boston Consulting Group, Embarcadero Center Suite 2400, San Francisco, CA 94111

**Keywords:** Splicing, U4/U6, Brr2, RNA helicase, ATPase, spliceosome activation

## Abstract

Before the spliceosome engages a pre-mRNA to excise its introns, the catalytic small nuclear RNA (snRNA) U6 is inactive because of base pairing with U4 snRNA; thus, spliceosome activation requires unwinding of base paired U4/U6, composed of stem I and stem II. The Ski2-like ATPase and RNA helicase Brr2p facilitates U4/U6 unwinding and the ultimately irreversible release of U4; however, the molecular mechanism behind Brr2p-mediated U4/U6 unwinding and the roles of the snRNAs in unwinding remains incompletely understood. To investigate the mechanism in vivo in budding yeast, we screened an unwinding deficient, cold-sensitive *brr2* mutant, associated with retinitis pigmentosa in humans, for genetic interactions with mutations in U4 snRNA. Destabilizing U4 mutations in either stem I or stem II suppressed the *brr2* mutant, providing functional evidence that Brr2p disrupts both stems in vivo. Further, destabilizing mutations in the intervening 5’ stem loop of U4 also suppressed the *brr2* mutant, and in vitro Brr2p displaced Prp31p from this stem loop, implicating Brr2p in disruption of this structure, too. Unexpectedly and counterintuitively, many destabilizing mutations in U4/U6 stem I exacerbated the *brr2* mutant. These mutations disrupted an intramolecular stem loop (U4-ISL1) in U4 that is mutually exclusive with U4/U6 stem I. We found that U4-ISL1 is required for splicing in vivo and for U4/U6 unwinding in vitro. Altogether, these results implicate Brr2p in disrupting all U4 secondary structures upstream of its initial U4 binding site and implicate an important role for U4 in antagonizing U4/U6 reannealing during Brr2p-mediated U4/U6 unwinding.

## Introduction

Splicing of nuclear pre-mRNA is a ubiquitous process in eukaryotes required for the generation of mature mRNA. Splicing is performed by a large ribonucleoprotein (RNP) complex termed the spliceosome, which catalyzes two consecutive transesterifications to remove an intron and ligate the two flanking exons (Wilkinson et al. 2020). First, a 2’ hydroxyl group from a conserved intronic adenosine, within the so-called branch point (BP) sequence, attacks the phosphodiester bond at the 5’ splice site (SS), generating a free 5’ exon and a lariat intermediate. Second, the 3’ hydroxyl group of the 5’ exon attacks the phosphodiester bond at the 3’ SS generating a mature mRNA and a lariat intron (Will and Lührmann 2011). The spliceosome is composed of five small nuclear RNP complexes (snRNPs), assembled on U1, U2, U4, U5 and U6 small nuclear RNAs (snRNAs), and numerous conserved non-snRNP proteins. The U6 snRNA catalyzes the two steps of splicing by adopting a conformation that coordinates catalytic divalent cations (Fica et al. 2013; Wan et al. 2019; Fica et al. 2017), in a manner similar to self-splicing group II introns (Toor et al. 2008). However, before binding to a splicing substrate, this catalytic conformation of U6 is precluded by extensive base pairing to U4 through two regions called stem I and stem II in a U4/U6-U5 triple snRNP (tri-snRNP), which recruits U4, U6, and U5 to a splicing substrate (Wilkinson et al. 2020). Activation of the spliceosome therefore requires separation of U6 from U4. Despite the critical importance of U4/U6 unwinding, the molecular mechanism of this reaction remains incompletely understood.

Formation of an active spliceosome starts with recognition of the 5’ SS by U1 and the BP by the BP binding protein, which is subsequently replaced by U2 after which a tri-snRNP complex joins, resulting in a pre-B complex that contains all five snRNAs (Charenton et al. 2019). A dramatic structural rearrangement follows where U1 and U4 are released, and the spliceosome is activated for catalysis.

The release of U1 from the 5’ SS requires the DEAD-box ATPase Prp28p (Charenton et al. 2019; Staley and Guthrie 1999), enabling the 5’ SS to base pair with the ACAGAGA sequence of U6 and to define the site of cleavage (Bertram et al. 2017; Lesser and Guthrie 1993; Kandels-Lewis and Séraphin 1993). The subsequent release of U4 from U6, the central rearrangement during spliceosomal activation, requires the Ski2-like ATPase Brr2p (Laggerbauer et al. 1998; Kuhn and Brow 2000; Kim and Rossi 1999; Raghunathan and Guthrie 1998; Noble and Guthrie 1996; Xu et al. 1996). Despite the expected requirement for Brr2p in all tissues, mutations in *BRR2* that compromise U4/U6 unwinding have been implicated in a human eye disease, retinitis pigmentosa (Zhao et al. 2009). After U4 is released several proteins including the nineteen complex (NTC) bind the spliceosome preparing it for the first catalytic step. After splicing has been completed and the mature mRNA released, recycling of the factors is required. For example, Prp24p must reanneal U6 to U4 to form the U4/U6 di-snRNP after which the U5 snRNP binds, reforming the tri-snRNP (Didychuk et al. 2018; Shannon and Guthrie 1991; Raghunathan 1998).

In *S. cerevisiae* splicing requires eight RNA-dependent ATPases that are conserved across eukaryotes (Cordin and Beggs 2013); seven of these ATPases transiently associate with the spliceosome, while Brr2p is an integral part of the U5 snRNP and therefore remains associated with the spliceosome from tri-snRNP recruitment through to spliceosome disassembly. Several ATPases function by a pulling mechanism (Semlow et al. 2016), disrupting their target at a distance and have accordingly been detected at the periphery of the spliceosome (Galej et al. 2016; Fica et al. 2017). Consistent with membership in the Ski2-like family of superfamily 2 (SF2) helicases, Brr2p-mediated RNA duplex unwinding requires a 3’ overhang and unwinds with a 3’ to 5’ polarity (Fairman-Williams et al. 2010).

The persistent association of Brr2p with the spliceosome necessitates regulation to avoid premature unwinding. The molecular mechanism behind Brr2p activation remains under investigation, but formation of the U6-5’SS interaction has been implicated in triggering a cascade of rearrangements that lead to movement of Brr2p from a position where it cannot bind U4 to a position where Brr2p directly interacts with U4 downstream of U4/U6 stem I (Kuhn and Brow 2000; Brow 2019; Charenton et al. 2019; Zhang et al. 2024). Because Brr2p is a 3’ to 5’ helicase, this placement of Brr2p on U4 strongly suggests that Brr2p unwinds U4/U6 stem I (Figure 1A)(Plaschka et al. 2017) in agreement with in vitro unwinding experiments (Mozaffari-Jovin et al. 2012). Indeed, in vivo cross-linking experiments between Brr2p and U4 demonstrated cross-links to the stem I region (Figure 1A)(Hahn et al. 2012). However, there is no functional evidence yet that Brr2p unwinds stem I in vivo. Brr2p also cross-links to the 3’ side of the 5’ stem loop in U4 (5’ SL, Figure 1A)(Hahn et al. 2012), which separates stem I from stem II, suggesting that Brr2p unwinds the 5’ SL, in addition to stem I. The U4 5’ SL is tightly bound by Snu13p and Prp31p, which would be expected to dissociate if Brr2 unwound the 5’ SL; however, in vitro unwinding of a minimal U4/U6 snRNP complex suggested that Snu13p and Prp31p remain bound to U4 (Theuser et al. 2016), raising questions as to the fate of the 5’SL and associated proteins during U4/U6 unwinding. Lastly, Brr2p does not cross-link to the U4 strand of stem II, raising questions as to how stem II is unwound (Figure 1A)(Hahn et al. 2012).

**Figure 1.**
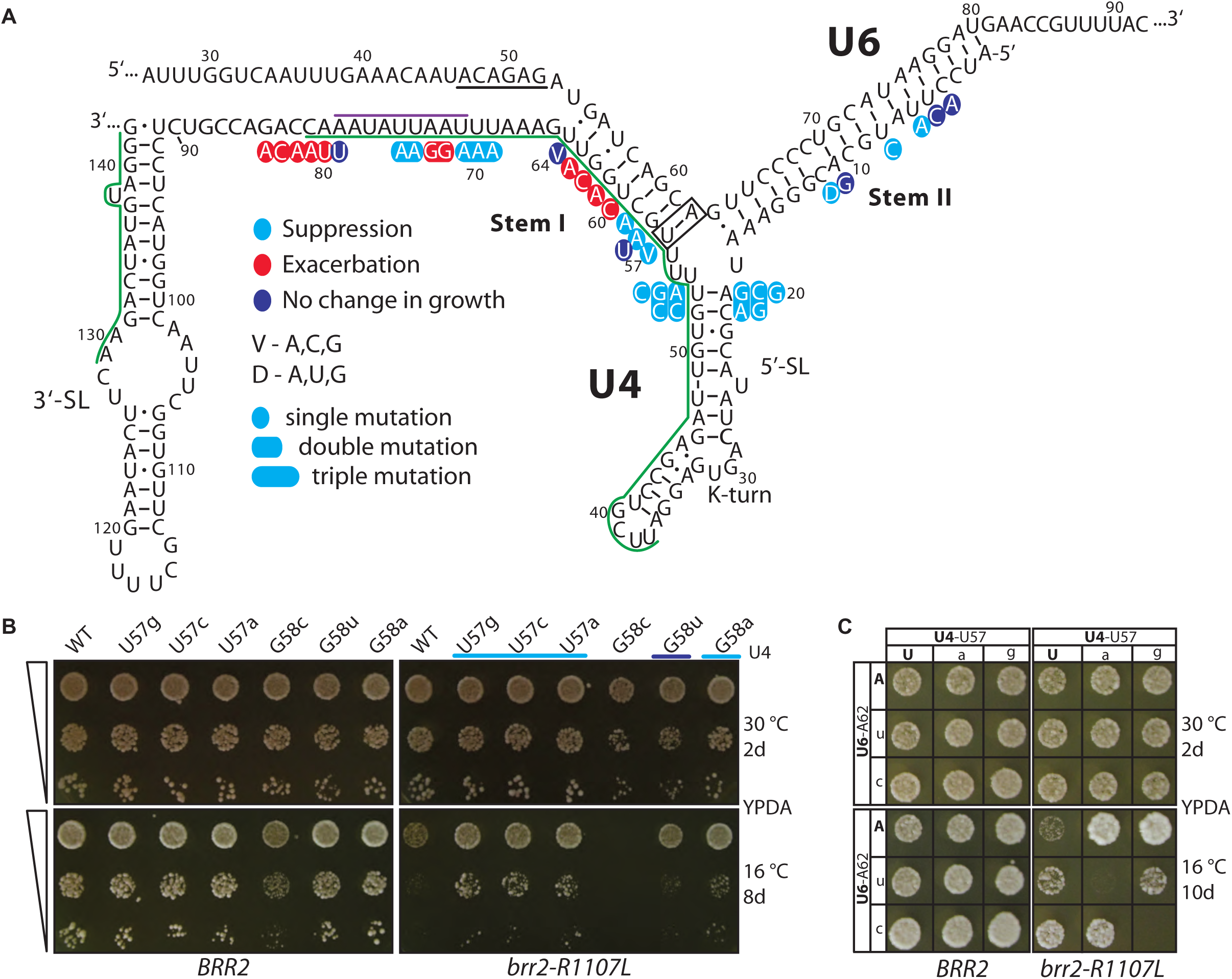
Evidence Brr2p unwinds U4/U6 stem I in vivo. (A) Schematic of base-paired U4/U6. Stem I and stem II are labelled. U4 mutations and their phenotypes in a *brr2-R1107L* mutant background are indicated and color coded for their phenotypes, as noted in the key. The U6-ACAGAG sequence is underlined, and the base pair investigated in (C) is boxed in black. The position of Brr2p on U4 within the preB-complex is indicated by a purple line. The in vivo crosslinking between Brr2p and U4 is indicated by a green line. (B) Destabilizing mutations in U4 stem I at positions G57 and G58A suppress the cold sensitive growth of *brr2-R1107L*. A dilution series of cultures (triangles) of the indicated yeast strains were frogged onto YPDA media and grown as indicated. The phenotype of each mutation is color coded, as in panel A, below the mutation; G58C is complicated by its cs phenotype in wild type *BRR2*, which correlates with a U4/U6 annealing defect (Figure Supplementary 2). (C) A compensatory analysis of U4/U6 base pair U57/A62 demonstrates that mutations in either the U4 or U6 strand of stem I suppress *brr2-R1107L* by disrupting base pairing. The indicated yeast strains were grown on YPDA media as indicated. The identity of the U4 or U6 alleles are shown in each column or row, respectively. Wild-type alleles are capitalized and in bold; mutants are lower case. Watson–Crick combinations fall on the diagonal (upper left to lower right).

After unwinding of U4/U6 stem I, U6 nucleotides released from stem I ultimately interact with U2 forming U2/U6 helix Ia, which juxtaposes the 5’SS and BP for 5’SS cleavage (Madhani and Guthrie 1992; Sun and Manley 1995; Mefford and Staley 2009) and helix Ib, which includes the conserved AGC triad that is involved in base triple interactions important for configuring the active site for coordination of catalytic divalent cations (Yan et al. 2016; Fica et al. 2014). After unwinding of U4/U6 stem II, the released U6 nucleotides ultimately form an intra-molecular stem loop called the U6-ISL, which participates in the catalytic base triples and catalytic metal ion positioning (Fica et al. 2013; Hang et al. 2015). Interestingly, psoralen cross-linking in the minor spliceosome (Frilander and Steitz 2001) revealed an intermediate in which U4/U6 stem I was disrupted and U2/U6 helix I was formed but U4/U6 stem II remained intact, implying that unwinding of stem I and stem II are temporally discrete events.

In order to investigate the mechanism of Brr2p-mediated release of U4 from U6 in vivo, we genetically and biochemically dissected the role of U4. We identified genetic interactions between *brr2-R1107L* and both stem I and stem II, consistent with unwinding of both stem I and stem II by Brr2p. Further, genetic and biochemical results support Brr2p-mediated unwinding of the intervening 5’SL with a concomitant release of Prp31p. Together, these observations support a role for Brr2p in destabilizing all RNA structures upstream from where Brr2p initially engages U4. Surprisingly, by genetic and biochemical approaches, we further implicated an intra-molecular stem loop in U4 (U4-ISL1) in promoting unwinding of stem I, implying U4 is actively involved in its own release from U6. These data indicate that the ultimately irreversible release of U4 from U6 is challenged by reversible unwinding of U4/U6 stem I.

## Results

### Brr2p promotes unwinding of U4/U6 stem I

To investigate Brr2p unwinding of U4/U6 in vivo we employed a genetic approach. Mutations in helicases often lead to cold-sensitive (cs) phenotypes, because a compromised helicase struggles to unwind a target duplex at low temperature due to relative stabilization of the duplex target at low temperature. Thus, we would expect a cs *brr2* mutant to unwind its target inefficiently, and we would further expect destabilizing mutations in the target duplex to suppress the cs phenotype of a *brr2* mutant.

To screen for such suppressors, we chose the cs *brr2* mutation R1107L, which is based on a mutation in human BRR2/SNRNP200 (R1090L) that leads to retinitis pigmentosa (Zhao et al. 2009). This yeast mutant compromises unwinding in vitro and limits growth at low temperature. To test for evidence that Brr2p unwinds U4/U6 stem I in vivo, we introduced mutations that disrupt Watson-Crick base pairing between U4 and U6 in stem I from position 57 to position 64 in U4 snRNA (Figure 1A-B, 2B), which was expressed as the sole copy of *SNR14* from a low-copy plasmid in wild-type and *brr2-R1107L* strain backgrounds.

Consistent with a role for Brr2p in unwinding U4/U6 stem I, we found that the cs phenotype of *brr2-R1107L* was suppressed by mutations at position 57-59 (Figure 1A-C, Figure 2B), mutations we term class I mutations. If suppression by mutations, such as those at position U57, was due to unwinding of U4/U6 stem I per se, as opposed to an alternative structure, we would expect that mutations in U6 at position A62 would also suppress *brr2-R1107L,* and combining mutations in U4 and U6 that re-establish Watson-Crick base pairing would abolish suppression by single mutations in either strand. Indeed, mutations in U6 at position A62 to either C or U did suppress *brr2-R1107L*; further, combining these mutations with U4 mutations at position U57 that restored Watson-Crick base pairing abolished the suppression reciprocally (Figure 1C), demonstrating that the suppression was due to disruption of stem I, per se. The abolishment of suppression was allele-specific; none of the non-Watson-Crick base combinations abolished suppression, although these phenotypes were limited by the growth of the U6 mutations (Figure 1C). We conclude that Brr2p unwinds U4/U6 stem I in vivo, as anticipated.

**Figure 2.**
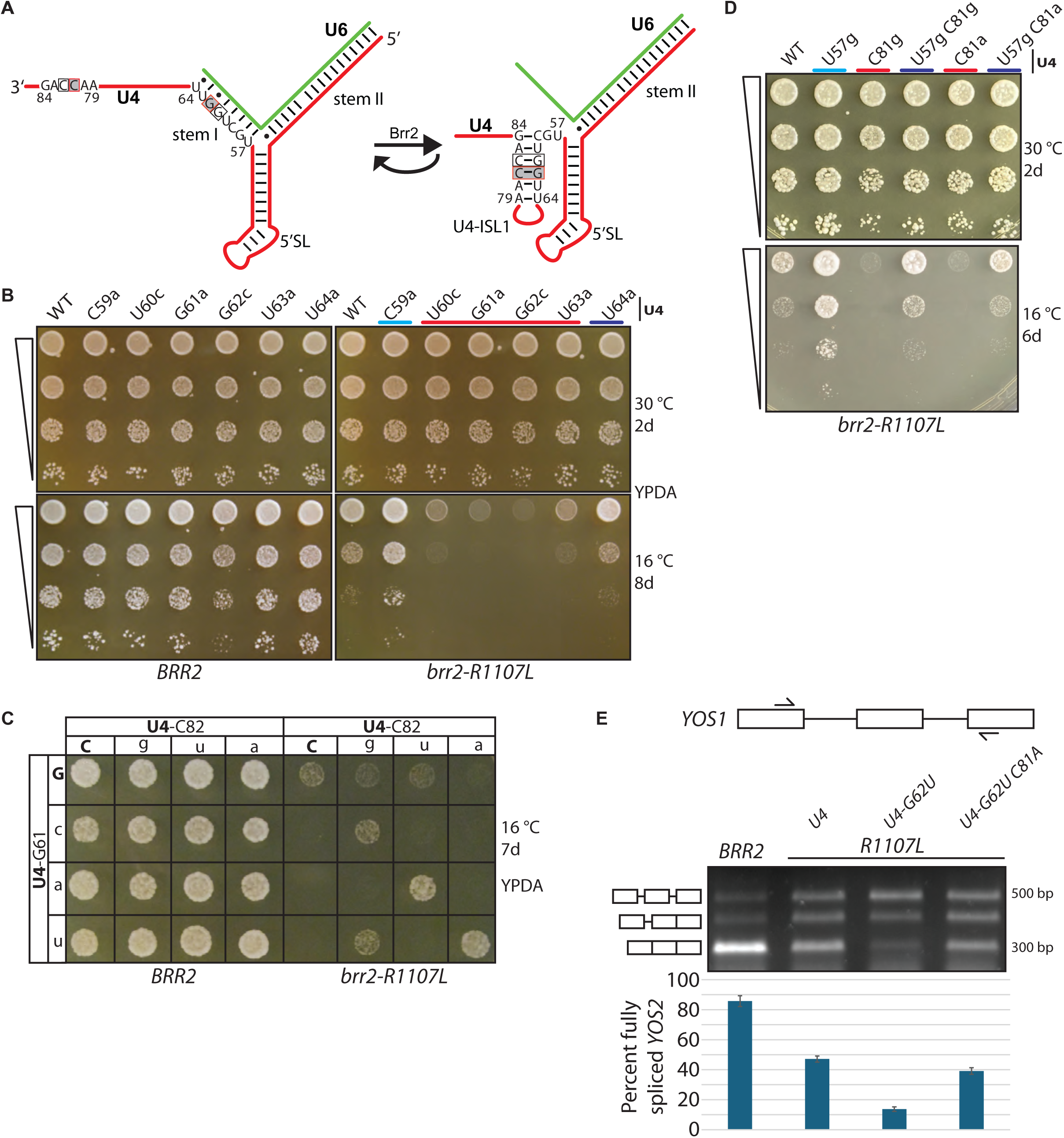
A novel intra-molecular U4 stem loop, mutually exclusive with U4/U6 stem I, cooperates with *BRR2*. (A) Schematic representation of U4/U6 indicating U4 nucleotides of stem I that after stem I disruption can pair with downstream residues to form an intra-molecular stem loop (U4-ISL1). The curved back-arrow depicts reversibility implied by the genetics. The base pair investigated in (C) is boxed in black and uncolored. The base pair and residue tested by RT-PCR in (E) is boxed in red and colored gray. (B) Mutations in U4 that destabilize stem I exacerbate *brr2-R1107L*. Growth was assessed and mutations were color-coded as in Figure 1B. (C) Compensatory analysis of U4-ISL1 base pair G61/C82 demonstrates that U4-ISL1 promotes *BRR2* function. Alleles are noted as in Figure 1C. Growth was assessed as in Figure 1C. (D) The stability of U4-ISL1 is balanced against the stability of U4/U6 stem I. Alleles are noted as in Figure 1B. Growth was assessed as in Figure 1B. (E) Compensatory analysis of U4-ISL1 base pair G62/C81 demonstrates that U4-ISL1 promotes splicing. Splicing of *YOS1* that contains two introns was analyzed by RT-PCR in the indicated strains. The schematic of *YOS1* shows the position of RT-PCR primers. The migration of the pre-mRNA, pre-mRNA with one intron spliced, and fully spliced mRNA is shown to the left of the agarose gel. Quantitation of 3 (U4-G62U) or 4 (the remaining strains) biological replicas with standard deviation (SD) is shown below. The quantification reflects the apparent splicing efficiencies since shorter species will be overrepresented. Yeast was grown to early log phase at permissive temperature of 30 °C and shifted to non-permissive temperature of 16 °C for 3 hrs.

### A stem loop in U4, ISL1, which is mutually exclusive with U4/U6 stem I, promotes Brr2-mediated unwinding of U4/U6 stem I

Surprisingly, we found that U4/U6 stem I mutations at positions 60-63 of U4 did not suppress the cold-sensitive growth phenotype of *brr2-R1107L* but instead exacerbated the growth phenotype (Figure 1A, Figure 2B); we term these mutations class 2 mutations. In principle, this class of mutations could reduce U4/U6 annealing and thereby compromise tri-snRNP formation and in turn synergize with the *brr2* mutation (however, see contrary evidence below). Alternatively, this class of mutations could antagonize Brr2p-mediated unwinding of U4/U6 stem I by preferentially destabilizing the product of unwinding, thereby facilitating re-annealing of U4/U6 stem I. For example, the mutations could destabilize an RNA secondary structure in U4 that is mutually exclusive with U4/U6 stem I and forms after stem I unwinding. Indeed, phylogenetic and biochemical studies implicated an intramolecular U4 stem loop within free U4 (Myslinski and Branlant 1991) that is mutually exclusive with U4/U6 stem I and that could serve to stabilize the unwound state of U4/U6 stem I and antagonize reannealing. With only minor adjustments, the stem of this structure includes the six U4 bases 79-84 on the 3’ side of the stem loop as well as the six U4 bases 59-64 on the 5’ side of the stem loop (Figure 2A). These 5’ residues all base pair in the stem loop in a manner mutually exclusive with their base pairing with U6 in U4/U6 stem I, so the mutations in these 5’ residues at positions 60-63 that exacerbated the *brr2* mutant disrupt not only U4/U6 stem I but also this stem loop. We named this extended stem loop U4-ISL1 (Figure 2A).

If the U4-ISL1 indeed forms after unwinding of U4/U6 stem I, then we would expect i) mutations on the other, 3’ strand of U4-ISL1 that similarly disrupt Watson-Crick base pairing to also exacerbate *brr2-R1107L*, and ii) second-site mutations that specifically restore base pairing within U4-ISL1 to abolish exacerbation. As noted, mutations at position U4-G61 strongly exacerbated *brr2-R1107L* (Figure 2B-C), and, indeed, mutations on the other side of the stem, for example U4-C82a, also exacerbated *brr2-R1107L* (Figure 2C). Further, second-site, compensatory mutations at positions U4-G61 or U4-C82 that restored base pairing within U4-ISL1 restored growth of *brr2-R1107L* in all cases (Figure 2C). Specifically, we found that the exacerbation of *brr2-R1107L* by the three point mutations at U4-G61 were each alleviated by compensatory mutations at position U4-C82 that restored Watson-Crick base pairing and vice versa – i.e., alleviation is mutual in all three cases (Figure 2A, C). Formation of a wobble base pair also alleviated the exacerbation (Figure 2C). The restoration of growth was allele specific because U4 mutations that did not restore base pairing within U4-ISL1 did not restore growth (Figure 2A, C).

Similarly, just as mutations at positions U4-U60, U4-G62 and U4-U63 exacerbated *brr2-R1107L*, mutations on the other side of the stem at positions U4-A83, U4-C81 and U4-A80 exacerbated *brr2- R1107L*, respectively, and mutations that restored Watson-Crick base pairing or lead to a wobble base pair alleviated this exacerbation in an allele-specific manner (Supplementary Figure 1A-C). Importantly, mutations that destabilized U4-ISL1 also compromised growth of a wild-type *BRR2* strain (see below), demonstrating a functional importance of U4-ISL1 formation independent of the *brr2-R1107L* mutation. These data provide evidence that U4-ISL1 promotes Brr2p function, and we propose that U4-ISL1 forms after unwinding of U4/U6 stem I to minimize reannealing.

If U4-ISL1 formed after unwinding of U4/U6 stem I to minimize reannealing, then we would expect these two structures to be, in effect, in equilibrium with one another, with the equilibrium dictated by the relative stabilities of the two structures. We would then also expect destabilizing mutations in each structure to offset one another, so that the impact of the mutations would be additive, rather than epistatic. To test this hypothesis, we combined a class 1 mutant, U4-U57G, which strongly suppressed *brr2-R1107L* (Figure 1B), with a mutant in the 3’ side of U4-ISL1 – either U4-C81G or -C81A, which both strongly enhanced *brr2-R1107L* (Supplementary Figure 1B), without also directly impacting U4/U6 stem I stability. In the context of the *brr2-R1107L* mutation, the double U4 mutants did indeed grow with an intermediate phenotype, relative to the individual U4 mutations; in fact, the double U4 mutants grew similar to wild type U4 in both cases, demonstrating that a productive balance between U4/U6 stem I and U4-ISL1 can be restored by destabilizing both structures (Figure 2D). These data support our hypothesis that U4/U6 stem I and U4-ISL1 are in equilibrium with one another and that U4-ISL1 serves to minimize reannealing of stem I and thereby to favor Brr2p-driven U4/U6 unwinding (see Discussion).

Notably some combinations of U4 ISL1 mutations that restored Watson-Crick base pairing grew better than the wild type U4 control. For example, for positions U4-G61 and U4-C82, mutations that gave rise to a U-A or A-U base pair in ISL1 grew better than wild-type U4 (Figure 2C). Because the U4-G61 mutations also destabilize U4/U6 stem I, we infer that these double U4 mutations suppressed *brr2* by the same mechanism that point mutations did at U4 positions 57-59 – that is, by destabilizing U4/U6 stem I but in this case without disrupting the U4-ISL1. These observations provide further evidence in vivo that Brr2p unwinds U4/U6 stem I.

To confirm the growth differences observed in the U4-ISL compensatory analyses reflected differences in splicing, we shifted yeast cultures to 16 °C for 3 hrs, harvested whole-cell RNA, and targeted *YOS1* transcripts for RT-PCR analysis (Figure 2E). In the wild-type *BRR2* strain, fully spliced mRNA represented more than 80% of all species; by contrast, in the *brr2-R1107L* mutant strain, fully spliced mRNA represented only ∼50% of all species (Figure 2E). Importantly, the mutation G62U, which disrupts the U4-ISL and exacerbates the *brr2-R1107L* growth defect, exacerbated the *brr2-R1107L* splicing defect, reducing fully spliced mRNA to ∼13%, and the compensatory mutation, C81A, which restored base pairing with position 62 in the U4-ISL1, restored fully spliced mRNA to ∼40% (Figure 2E). Thus, disrupting U4-ISL1 exacerbates the growth defect of the *brr2* mutant, because disrupting U4-ISL1 exacerbates the splicing defect of the *brr2* mutant.

Mutations in U4 snRNA that destabilized U4/U6 stem I could in principle affect U4/U6 annealing during tri-snRNP assembly in addition to or instead of affecting U4/U6 unwinding, especially for the class 2 mutants, which exacerbated *brr2*. In a wild-type strain all available U4 is base paired with U6 and no free U4 is detected. We could therefore test for defects in U4/U6 annealing by assaying for the accumulation of free U4 snRNA in mutants. We examined eight class 1 or class 2 mutants within U4/U6 stem I in the *brr2-R1107L* mutant background. For all mutants examined, we did not detect any free U4, except for one of the three class 1 mutants, U4-G58A, and U4-G58C, which conferred a growth defect in wild-type *BRR2* and was not examined further (Figure 1B; Supplementary figure 2A, C). Although U4-G58A showed evidence of a U4/U6 annealing deficiency, as a class 1 mutation, it nevertheless suppressed *brr2-R1107L*, which is limited for growth by U4/U6 unwinding, so we infer that U4-G58A suppressed by facilitating unwinding. In all other cases, we conclude that the U4 mutations in U4/U6 stem I do not compromise U4/U6 stem I annealing but rather either facilitate U4/U6 stem 1 unwinding (class 1 mutations) or compromise U4-ISL1 formation (class 2 mutations), after U4/U6 stem I unwinding.

### U4-ISL1 promotes U4/U6 unwinding in vitro

To test the importance of U4-ISL1 in a wild-type *BRR2* background, we disrupted ISL1 with two consecutive mutations in U4 at positions 61 and 62, changing GG to CC (abbreviated U4-GG61CC), or two consecutive mutations in U4 at positions 81 and 82, U4-CC81GG. Further, we combined these mutations to restore base pairing within U4-ISL1. Tellingly, both double mutations on their own conferred a cold-sensitive phenotype (Figure 3A) and combining the two double mutations, restored WT growth (Figure 3A), demonstrating that U4-ISL1 is important for growth even in wild-type cells. This cold-sensitive phenotype of the double mutations is not easily rationalized by a role for U4-ILS1 in U4/U6 annealing during tri-snRNP formation, given the mutual exclusivity with U4/U6 stem I and the expectation that disruption of U4-ISL1 would promote, not antagonize, annealing. However, the phenotype is readily explained by a role for U4-ILS1 in U4/U6 unwinding by Brr2p, particularly given that the cold-sensitivity of these mutations phenocopies the cold-sensitivity of the *brr2-R1107L* mutation. To test directly whether these U4-ISL1 mutations impact U4/U6 unwinding, rather than U4/U6 annealing in a wild-type *BRR2* context, we uncoupled U4/U6 unwinding from U4/U6 annealing by isolating U4/U6-bound tri-snRNPs by pulldown from budding yeast extract; this isolation excludes the U4/U6 annealing factor Prp24p (Raghunathan and Guthrie 1998). To test unwinding of U4/U6 in vitro within tri-snRNPs assembled in vivo with mutated U4, we grew strains expressing TAP-tagged *BRR2* and the U4 ISL1 mutations that compromised growth at low temperature in a wild-type *BRR2* strain (Figure 3A). As with the growth assay, we compared the two double mutations GG61CC and CC81GG alone and in combination. We verified the integrity of tri-snRNPs by primer extension (Figure 3B). Relative to U4, we observed a ∼2-fold enrichment of U5 in both the GG61CC and CC81GG strains, indicating an increase in Brr2p-bound free U5 snRNPs relative to tri-snRNPs, suggesting a modest defect in tri-snRNP assembly that may indicate U4/U6 annealing is partially compromised by the mutations; importantly, the pre-annealed U4/U6 in the isolated tri-snRNP allows us to bypass such defects when investigating U4/U6 unwinding.

**Figure 3.**
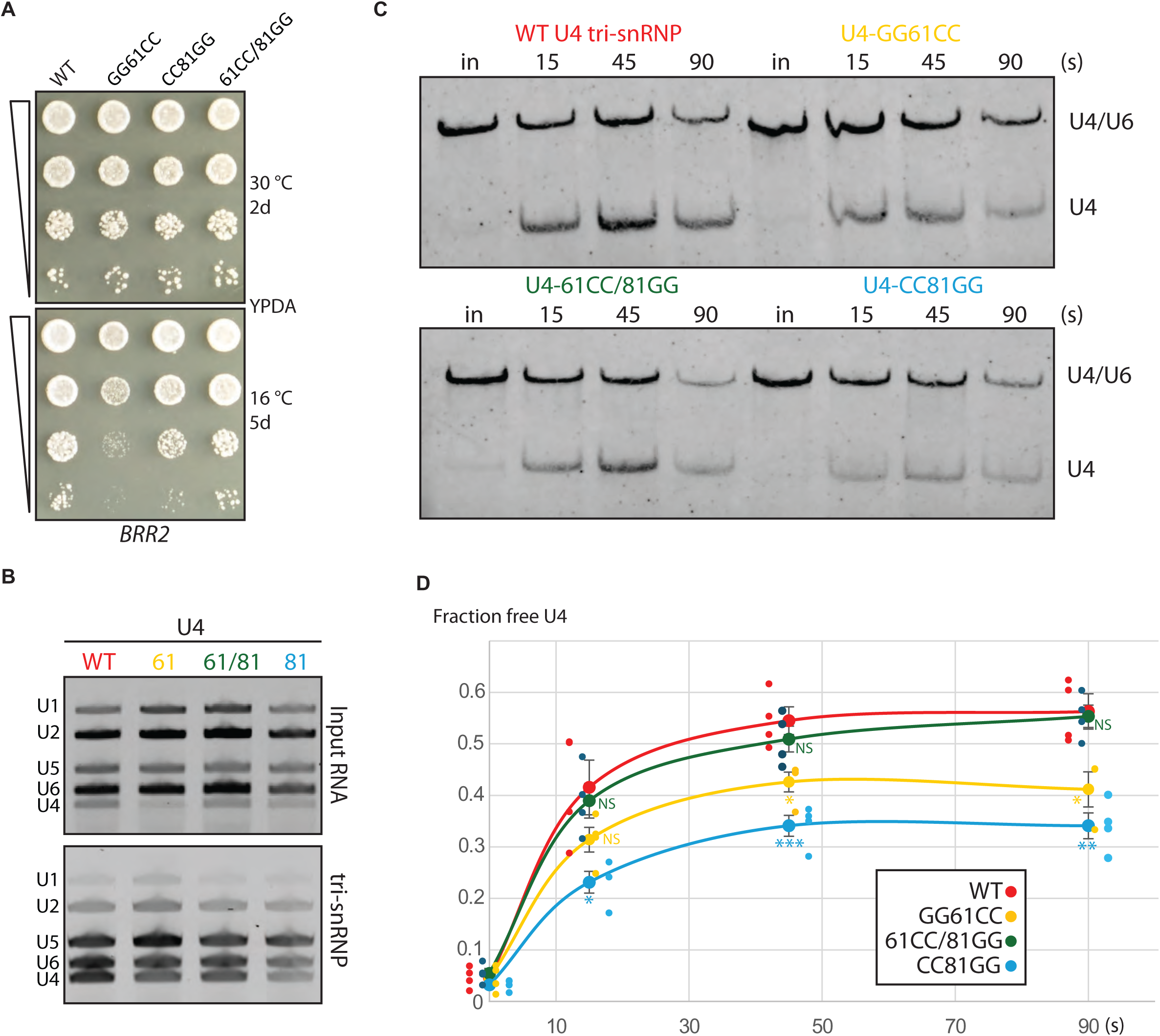
U4-ISL1 promotes U4/U6 unwinding in vitro, in the context of tri-snRNPs. (A) U4-ISL1 is required for growth in a wild-type *BRR2* strain. Simultaneous disruption of G61/C82 and G62/C81 ISL1 base pairs with the double mutations GG61CC or CC81GG compromised growth; combining these mutations restored base pairing and growth. Growth was assessed as in Figure 1B. (B) Isolation of wild-type and U4-ISL1 mutant tri-snRNPs, via pull down of TAP-tagged Brr2p. Primer extension of extracted snRNA shows expected enrichment of U4, U5, and U6, relative to input for the pull down. (C) Unwinding of U4/U6 in tri-snRNPs, isolated from a wild-type strain or a U4 mutant strain in which U4-ISL1 is destabilized. Unwinding was assayed at the indicated timepoints. Base paired U4/U6 and free U4 were monitored on non-denaturing gel by northern blotting for U4. “In” stands for “input”. (D) Quantitation of U4/U6 unwinding in panel C shows U4-ISL1 is required for efficient U4/U6 unwinding. Fraction of free U4 is shown as a function of time for four separate experiments. A p-value range is indicated for the three timepoints for the mutant versus the wild-type tri-snRNPs, based on an unpaired T test score (two-tailed), with * indicating p<0.05; **, p <0.01, ***, p<0.001; and NS, not significant; standard error of the mean is also shown. In panels B-D, the tri-snRNPs are color coded, according to their identities as indicated.

We initiated unwinding of U4/U6 in the isolated tri-snRNPs by adding ATP and sampled time points for unwound U4/U6 (Figure 3C). Quantification of free U4 (Figure 3D) revealed reduced unwinding for both double mutants, whereas the quadruple mutant that restored complementarity in U4-ISL1 also restored unwinding to levels indistinguishable from the wild-type U4 (Figure 3C-D). These results demonstrate that the stability of U4-ISL1 positively correlates with U4/U6 unwinding in tri-snRNPs.

### U4-ISL1 is semi-conserved between lower and higher eukaryotes

A structure named hairpin II that is related to U4-ISL1 was previously proposed to form across eukaryotes in free U4 (Myslinski and Branlant 1991). To investigate whether the potential to form either hairpin II or U4-ISL1 is conserved, we inspected an alignment of the 177 seed sequences from the RFam U4 family (Ontiveros-Palacios et al. 2024). First, we inspected relatives of *S. cerevisiae* belonging to a subset of fungi associated with a single node of a U4 phylogenetic tree derived from these seed sequences. Though related, by the Jukes-Cantor model (Jukes and Cantor 1969), the U4 sequences of these fungi have diverged from *S. cerevisiae* (distance≤0.11; Figure Supplementary 3A, middle row) to a degree that is comparable to the divergence of U4 sequence from humans to fish (distance≤0.07; Figure Supplementary 3A, bottom row). Notably, all fungi seed sequences associated with this node preserve the ability to form U4-ISL1 (Figure Supplementary 3B). A difference between *S. cerevisiae* and *H. sapiens* in the conserved U4 strand of U4/U6 stem I is that a GG sequence in *S. cerevisiae* (positions 61-62) has transitioned to an AA sequence in *H. sapiens* (Figure Supplementary 3B, in bold). While this change disrupts U4-ISL1, a four nucleotide shift in the register of the downstream strand allows the formation of an alternative ISL – hairpin II (Myslinski and Branlant 1991). Importantly, hairpin II, like U4-ISL1, is mutually exclusive with U4/U6 stem I (Figure Supplementary 3B). To compare the secondary structures of unwound U4 from either human or yeast (Reuter and Mathews 2010), we focused on the first 90 bases of U4, which includes the 5’-SL and the single-stranded RNA between the 5’-SL and the 3’-SL but not the 3’SL itself (for *S. cerevisiae* see figure 1A), because U4 alignments from RFam demonstrates that the 3’ end of U4 is much more divergent than the 5’ end (Ontiveros-Palacios et al. 2024). The “RNAstructure” software package predicted two structures for *S. cerevisiae* both of which included the six base-pair U4-ISL1 and one of which included an elongated 5’-SL (Figure Supplementary 3A, left side). For *H. sapiens* two structures were predicted one of which also included an elongated 5’-SL as well as the previously predicted four base-pair hairpin II (Figure Supplementary 3A, right side) (Myslinski and Branlant 1991). In both species, the intramolecular stem loop formed just downstream of the extended 5’-SL, suggesting these secondary structures might stack on one another. These similarities suggest that while the stem loop is shifted between yeast and humans, the stem loop is conserved between these species. Indeed, an alignment of U4 sequences demonstrates that all analyzed species, excluding the fungi node, have the potential to form hairpin II (Figure Supplementary 3B). Altogether, these results suggest that two similar hairpins, U4-ISL1 and hairpin II, serve the same function across eukaryotes – to prevent reannealing of U4/U6 stem I, after Brr2p-mediated unwinding.

### A second stem loop, U4-ISL2, impedes Brr2p activity

In determining if U4-ISL1 is essential for growth, we also introduced mutations in U4 at positions 79-84 of the 3’ strand of the stem that would completely destabilize U4-ISL1 (Figure 4A), without impacting U4/U6 stem I residues. To our surprise, compared to single mutations that disrupted U4-ISL1 and exacerbated *brr2-R1107L* (e.g., U4-C81A), the hextuple 79-84 mutant and related mutants exacerbated *brr2-R1107L* less strongly (Figure 4B, compare column 3-5 to column 2). Careful examination of the U4 sequence for competing structures suggested a second ISL, we termed ISL2, that could form between nucleotides 79-85 (that includes the 3’ strand of ISL1) and 69-75 (in the loop of ISL1) by virtue of four Watson-Crick base pairs and one wobble base pair (Figure 4A). Whereas U4-ISL2 would be mutually exclusive with U4-ISL1, ISL2 would be compatible with U4/U6 stem I and consequently could form at an earlier stage than ISL1. Indeed, given that ISL2 occludes the Brr2p binding site in U4 from which U4/U6 unwinding is thought to initiate (Hahn et al. 2012; Charenton et al. 2019; Mozaffari-Jovin et al. 2012; Zhan et al. 2018; Bai et al. 2018), U4-ISL2 could potentially antagonize Brr2p engagement of U4 and consequently U4/U6 stem I unwinding. Such an impact would rationalize why greater disruption of ISL1 appeared to mitigate exacerbation of the *brr2* mutant by milder disruption of ISL1 – that is, the more extensive hextuple mutant also disrupted U4-ISL2, thereby facilitating recognition of U4 by Brr2p for unwinding.

**Figure 4.**
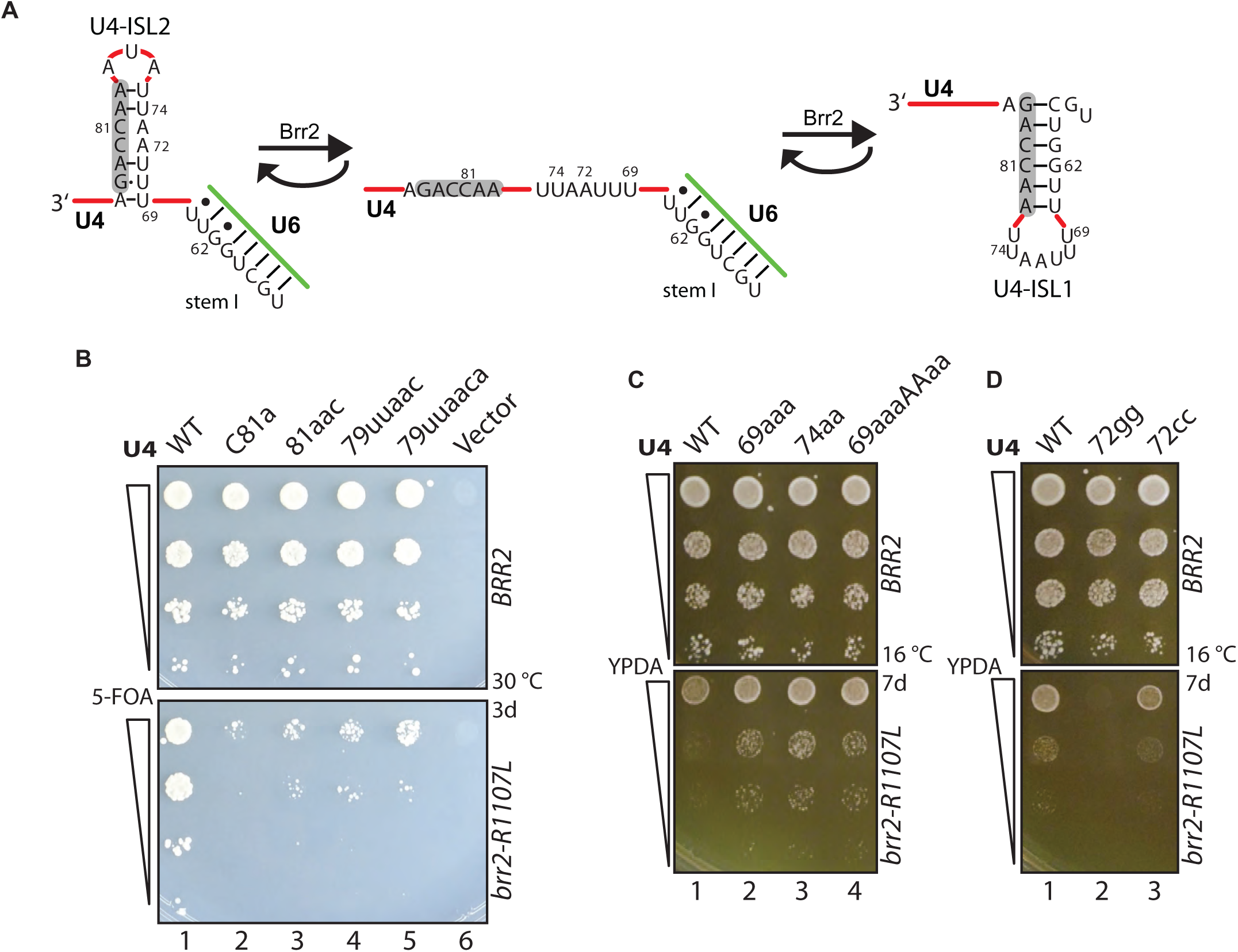
A second, intra-molecular U4 stem loop (U4-ISL2), which is mutually exclusive with U4-ISL1, antagonizes *BRR2.* (A) Schematic representation of U4/U6 showing U4 nucleotides that can form U4-ISL2, unwinding of U4-ISL2 to accommodate Brr2p binding, and formation of U4-ISL1 after Brr2p-mediated U4/U6 stem I unwinding. The U4 strand that participates in both U4-ISL1 and -ISL2 is highlighted in gray to illustrate the mutually exclusive nature of these stem loops. (B) Mutating three, five, or all six positions within the downstream side of the stem in U4-ISL1 has a less severe growth phenotype than a single point mutation, suggesting an impact on an alternative structure. Growth was assessed as in Figure 1B but on solid 5-FOA media. (C, D) Destabilizing U4-ISL2 suppress *brr2-R1107L* (C), whereas stabilizing ISL2 exacerbates *brr2-R1107L* (D). In both panels, the tested mutations alter the upstream strand of the stem in U4-ISL2, without perturbing U4-ISL1. In panel C, all three mutations destabilize U4-ISL2. In panel D, the 72gg mutation hyperstabilizes U4-ISL2, whereas the 72cc mutation does not. Growth was assessed as in Figure 1B.

To test explicitly whether U4-ISL2 modulates Brr2p activity, we introduced mutations within U4 residues 69-75, in the loop of U4-ISL1, that either destabilized U4-ISL2 (Figure 4C, columns 2-4) or stabilized U4-ISL2 (Figure 4D, column 2). In contrast with mutations in residues 59-64 that weakened U4-ISL1 and exacerbated *brr2-R1107L*, mutations that weakened U4-ISL2 suppressed *brr2-R1107L*; conversely, mutations that strengthened U4-ISL2 exacerbated *brr2-R1107L*, in an allele-specific manner (Figure 4C-D). Additionally, mutations in the bulge on the opposite, 3’ strand of ISL2, that neither weakened nor stabilized ISL2 but destabilized ISL1 (e.g., U4-C81A), exacerbated *brr2-R1107L*, as expected (Figure 2C-D, Figure 4B). Together, these data establish evidence that U4-ISL2 antagonizes Brr2p and provide the first functional evidence in vivo consistent with Brr2p binding to ssRNA in this region of U4.

### Brr2p promotes unwinding of U4/U6 stem II

Given our genetic evidence that Brr2p unwinds U4/U6 stem I, we tested for genetic evidence that Brr2p also unwinds U4/U6 stem II, especially given cross-linking data that suggests Brr2p only unwinds U4/U6 stem I and not stem II (Figure 1A)(Hahn et al. 2012). To investigate whether Brr2p unwinds U4/U6 stem II, we destabilized stem II by mutating eight positions in the U4 strand of stem II and tested for suppression of *brr2-R1107L*; we only investigated mutations that were not already cold-sensitive in a wild-type *BRR2* background (Hu et al. 1995). Mutations at three positions (position 6, 8 and 12) suppressed *brr2-R1107L*; mutations at the remaining positions were aphenotypic (Figure 5B).

**Figure 5.**
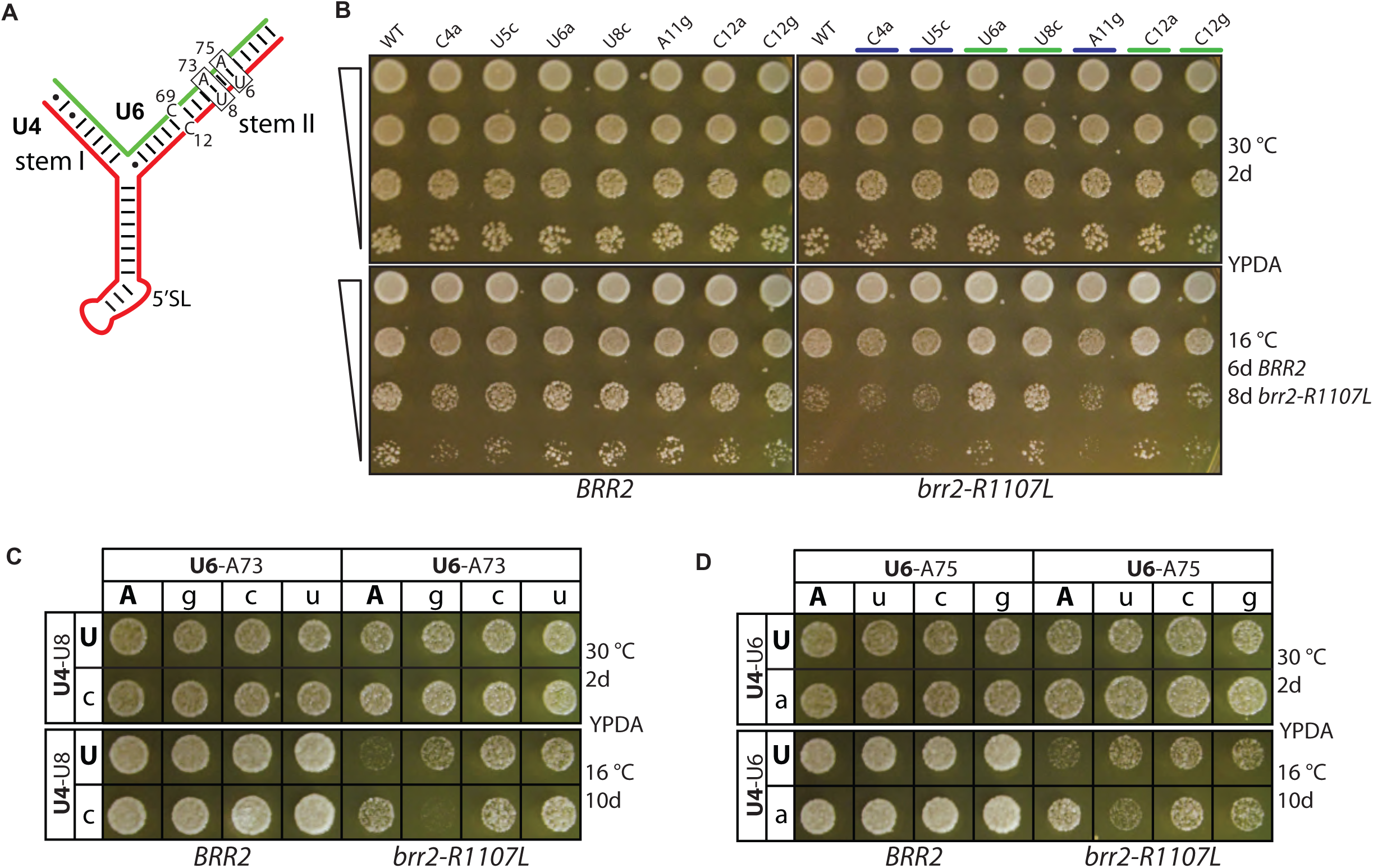
Evidence Brr2p unwinds U4/U6 stem II in vivo. (A) Schematic representation of U4/U6 stem II indicating U4 and U6 nucleotides where mutations suppress *brr2-R1107L*. (B) U4 mutations at positions 6, 8 and 12 within stem II suppress *brr2-R1107L*. Growth was assessed as in Figure 1B. (C, D) Compensatory analysis of U4/U6 base pairs U8/A73 (C) and U6/A75 (D) demonstrates that mutations in the U6 strand of stem II, in addition to the U4 strand, suppress *brr2-R1107L* and that mutations in both strands suppress by disrupting stem II base pairing. Growth was assessed as in Figure 1C.

If mutations at position 6 and 8 in U4 suppressed *brr2-R1107L* by destabilizing stem II, we would expect similar suppression of *brr2-R1107L* by point mutations in U6 that disrupt the same base pair.

Furthermore, combining mutations in U4 and U6 at these positions that restored base pairing should result in a loss of suppression. We indeed observed suppression of *brr2-R1107L* by point mutations at U6-A73, which is base paired to position U8 of U4 snRNA, and in U6-A75, which is base paired to position U6 of U4 snRNA (Figure 5A, C-D). Further, combining mutations U4-U8C and U6-A73G or mutations U4-U6A and U6-A75U, which restored base pairing in each case, indeed reciprocally abolished suppression and in an allele-specific manner (Figure 5A, C-D). These results indicate that mutations in both the U4 and U6 strands of stem II suppress *brr2-R1107L* and do so by disrupting U4/U6 stem II, implying that Brr2p is required to unwind stem II, in addition to stem I (see Discussion).

Residue U4-C12 is the only residue within U4/U6 stem II that does not form a base pair with U6; the opposite position in U6 is also a cytosine. Surprisingly, all three mutations of U4-C12 suppressed *brr2-R1107L*, including U4-C12G, which we expected to exacerbate *brr2* by hyperstabilizing stem U4/U6 stem II (Figure 5A and Supplementary Figure 4). Mutations of U6-C69, which are opposite U4-C12 in stem II, did not suppress *brr2-R1107L* (Figure Supplementary 5G-H). Notably, the U6-C69G mutation would also hyperstabilize stem II (as U4-C12G), which we expected to exacerbate *brr2*, but U6-C69G did not differentially affect *brr2-R1107L* compared to U6-C69A or -C69U (Figure Supplementary 5G-H), suggesting that position 12 of U4 and position 69 of U6 do not readily base pair. Human U4/U6 has a similar pyrimidine pair at position 13 of U4 in U4/U6 stem II (Brow and Guthrie 1988), suggesting a function for this bulge. Within the yeast precatalytic B-complex (Bai et al. 2018), U4-C12 is near Prp3p H308, which is conserved and causes retinitis pigmentosa when mutated in humans (Zhong et al. 2016), suggesting that the suppression of *brr2-R1107L* by mutations at position U4-C12 might be due to disruption of RNA-protein interactions that may indirectly destabilize U4/U6 stem II.

### No evidence that formation of either U2/U6 helix I or U6-ISL promotes U4/U6 unwinding

U6 nucleotides, released after unwinding U4/U6 stem I, base-pair with U2 to form U2/U6 helix I. To test whether this U2/U6 interaction could promote U4/U6 stem I unwinding in cooperation with U4 ISL1 by similarly stabilizing the product of unwinding, we introduced mutations that would either i) disfavor formation of U2/U6 helix I by directly destabilizing the helix (Figure Supplementary 5A-C) or ii) favor formation of U2/U6 helix I, by destabilizing the competing stem I within U2 (Figure Supplementary 5B,D-E). If formation of U2/U6 helix I promotes U4/U6 stem I unwinding, we would expect disfavoring formation of U2/U6 helix I to exacerbate *brr2-R1107L* and favoring formation of U2/U6 helix I to suppress *brr2-R1107L*. To destabilize helix Ia, we mutated position U6-G55 to a C; we included mutations in the U6 intramolecular stem loop (Figure Supplementary 5C) to maintain this structure. U6-G55C did not impact growth of *brr2-R1107L* (Figure Supplementary 5A). To enhance formation of helix Ib, we tested a single mutation, U2-C15G, and a double mutation, U2-CC14GG, which each destabilize U2 stem I and thereby favor U2/U6 helix Ib (Figure Supplementary 5B,D-E). The single U2-C14G mutation has been demonstrated to suppress a mutation within U2/U6 helix Ib in U6 (U6-C61A)(Hilliker and Staley 2004), but neither mutation tested impacted growth of *brr2-R1107L* (Figure Supplementary 5D). These data provide no evidence that the formation of U2/U6 helix I promotes unwinding of U4/U6 stem I.

U6 nucleotides, released after unwinding U4/U6 stem II, form the catalytic U6-ISL, which might promote U4/U6 stem II unwinding by stabilizing the product of unwinding. To test whether U6 ISL formation promotes U4/U6 stem II unwinding, we tested whether destabilization of U6-ISL exacerbated the cold-sensitivity phenotype of *brr2-R1107L*. We examined nine mutations at four positions within the U6-ISL, only one of which is also involved in U4/U6 base pairing (position U6-C67). At position U6-A82 neither of the mutations tested (U and C) showed a genetic interaction (Figure Supplementary 5B,D,G) and at position U6-C84 none of the three substitutions showed a genetic interaction with *brr2-R1107L* (Figure Supplementary 5B,F-G). Additionally, we tested a substitution at U6-C67 (U) and all three substitutions at U6-C69, which all failed to show a genetic interaction with *brr2-R1107L* (Figure Supplementary 5B,G-H). Because mutations at position 67 disrupted both U4/U6 stem II and the U6-ISL, the negative results at this position might be explained by counterbalancing effects, but such effects cannot explain the results at the other three positions. These data provide no evidence that formation of the U6-ISL promotes unwinding of U4/U6 stem II (see Discussion).

### Brr2p activity disrupts the 5’SL in U4

The genetic results above demonstrate that Brr2p activity is required for unwinding of U4/U6 stem II, in addition to stem I. In two simple models, Brr2p would unwind both stem I and stem II by either pulling of U4 or translocating along U4 processively from the site at which Brr2p engages U4, downstream of U4/U6 stem I, through to the 5’ end of U4 (see Discussion). Both models predict unwinding of the 5’SL of U4, in addition to stems I and II. To test for destabilization of the 5’SL of U4, we introduced mutations at the closing base pairs of the U4-5’SL (positions 20-21 and 52-53, see Figure 1A and Figure 6A) that should destabilize the U4-5’SL and compensatory mutations that should restore base-pairing. Overall, destabilizing mutations suppressed *brr2-R1107L* (Figure 6A, columns 2-3, 6-7, 9-12), whereas compensatory mutations abolished suppression (column 4) or even exacerbated the *brr2-R1107L* mutant (column 8), possibly due to hyperstabilization of the 5’SL. These results are consistent with disruption of the U4-5’SL by Brr2p and suggest that proteins bound to the U4-5’SL are displaced.

**Figure 6.**
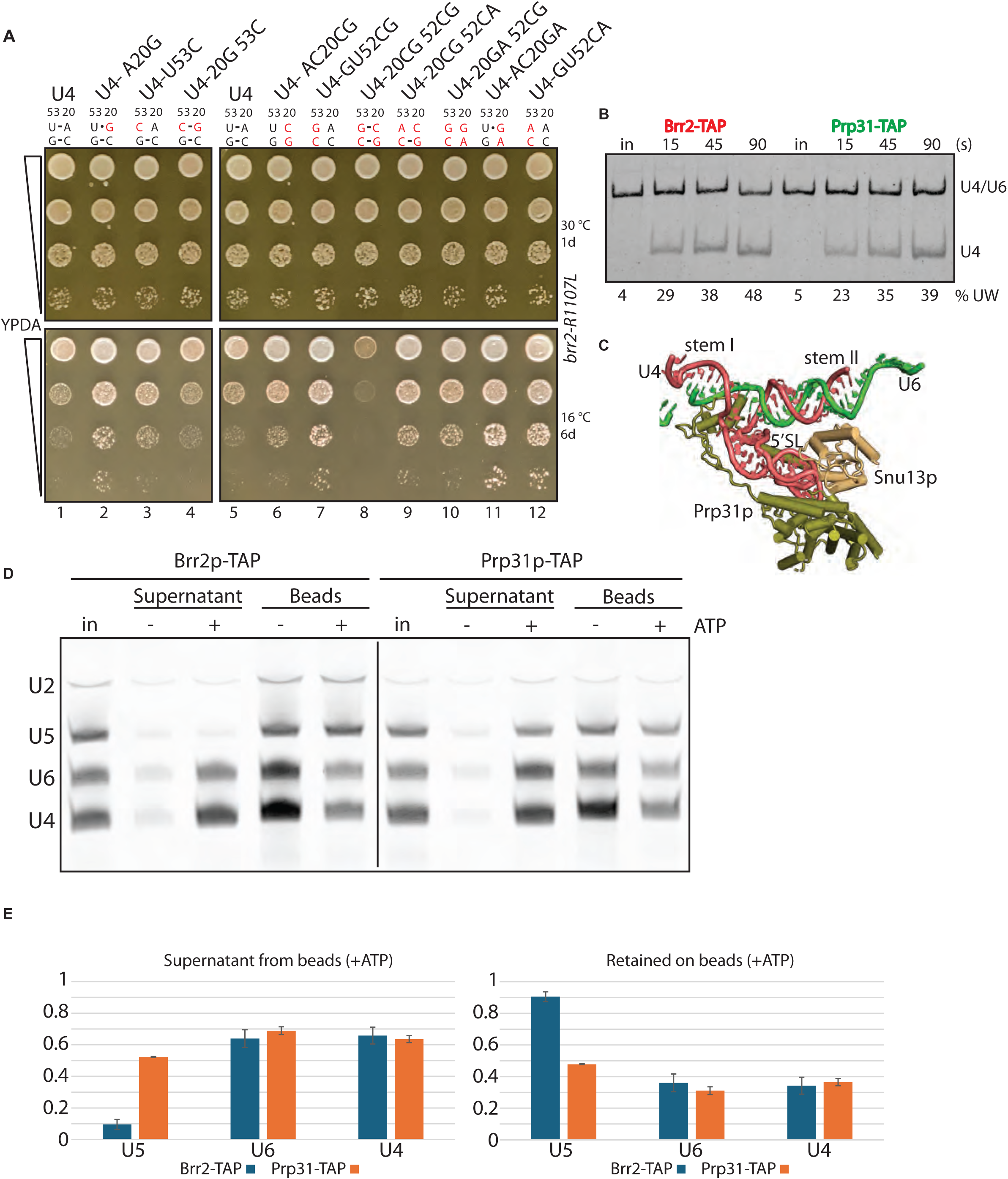
Evidence that Brr2p also unwinds the U4-5’SL. (A) Destabilization of the U4-5’SL suppresses *brr2-R1107L*. Single and double mutations in U4 that destabilize the U4-5’SL and suppress *brr2-R1107L* and compensatory mutations that restore base-pairing and abolish suppression are shown. The U4 allele names are indicated above each column, and the status of base pairs U53/A20 and G52/C21 of the U4-5’SL is shown just below; mutations are indicated in red. Growth was assessed as in Figure 1B. (B) Wild-type tri-snRNPs isolated via Brr2p-TAP or Prp31p-TAP unwind U4/U6 similarly. U4/U6 unwinding was assayed and displayed as in Figure 3C. Percent unwinding is displayed underneath the rows (% UW). (C) Overview of U4/U6 stem I and stem II and U4-5’SL with Snu13p and Prp31p bound. U4 is colored red and U6 is colored green. Generated using PyMOL, PDB 5NRL (Plaschka et al. 2017). (D) ATP-dependent unwinding releases Prp31p from U4 snRNA. The snRNAs associated with either IgG beads or released to the supernatant were separated after unwinding showing retention of U5 in Brr2p-TAP particles but release of U4 in Prp31p-TAP particles. (E) Panel D quantitation showing the fraction of U4, U5, or U6, that was released from (left) or retained on (right) the IgG beads, after incubation of Brr2p-TAP or Prp31p-TAP tri-snRNPs in the presence of ATP. Standard deviation is shown and n=3.

We consequently tested in vitro whether Brr2p displaces protein from the U4-5’SL. This stem loop is bound tightly by two proteins: Snu13p and Prp31p (Figure 6C)(Hardin et al. 2015). We assayed for dissociation of Prp31p from the U4-5’SL using tri-snRNPs bound to beads via TAP-tagged Prp31p; we expected that U4/U6 unwinding would release U5 and U6, and we assayed for release versus retention of U4 on beads. As a control, we compared tri-snRNPs bound to beads via TAP-tagged Brr2p; in this case we expected that U4/U6 unwinding would release U4 and U6 but retain U5, given Brr2p is an integral component of the U5 snRNP. The two tri-snRNPs assembled with similar efficiencies (Figure 6D, “in”) and the tri-snRNPs isolated on beads unwound U4/U6 with similar efficiencies (Figure 6B), indicating the Prp31p TAP tag did not perturb function. After unwinding by the addition of ATP, we assayed for snRNAs released into the supernatant or retained on beads (Figure 6D and quantified Figure 6E). As expected, with the TAP-tagged Brr2p snRNPs, ATP triggered efficient release of U4 and U6 (∼65%) but not U5 (∼10%, comparable to background release in the absence of ATP). With the TAP-tagged Prp31p snRNPs we observed similarly efficient release of U4 and U6 (∼65%) and also efficient release of U5 (∼50%), which may be slightly lower because of interactions between Prp31p and the core U5 snRNP protein Prp8p (Figure 6D-E). The efficient release of U4 from Prp31p supports disruption of the U4-5’SL by Brr2p, as implied by the genetic tests (Figure 6A). Note that competitor U4-5’SL RNA was required to observe efficient release of U4. In the absence of competitor only ∼30% of U4 was observed in the supernatant (Figure Supplementary 6), implying that Prp31p, after release from U4, readily rebinds the U4-5’SL/Snu13p, consistent with previous binding studies (Hardin et al. 2015). In conjunction with the data implying Brr2p disrupts U4/U6 stems I and stems II, these genetic and biochemical data imply that Brr2p disrupts all U4 structures upstream of the initial binding site on U4, consistent with processive pulling of or translocation along U4.

## Discussion

Unwinding of U4/U6, composed of stem I and stem II, is a critical step in spliceosome activation (Figure 1), but the mechanism remains incompletely defined. Even though Brr2p only cross-links in vivo to U4 in stem I and not in stem II (Hahn et al. 2012), we have found evidence that U4/U6 unwinding requires that Brr2p separate not only stem I but also stem II (Figures 1,5). Further, we found evidence that Brr2p destabilizes the intervening U4-5’SL and releases the bound protein Prp31p (Figure 6), consistent with cross-linking of Brr2p in vivo to the 3’ side of the U4-5’SL (Hahn et al. 2012). In addition, we discovered in vivo and in vitro that a stem loop in U4, ISL1 (Figure 2-3), which is mutually exclusive with U4/U6 stem I, surprisingly promotes Brr2p-mediated U4/U6 unwinding, likely by trapping the U4 strand of U4/U6 stem I after unwinding. Altogether, our data support a model in which Brr2p destabilizes all U4 secondary structures upstream of its initial binding site and in which U4 plays an active role in driving U4/U6 unwinding forward by preventing reannealing (Figure 7).

**Figure 7.**
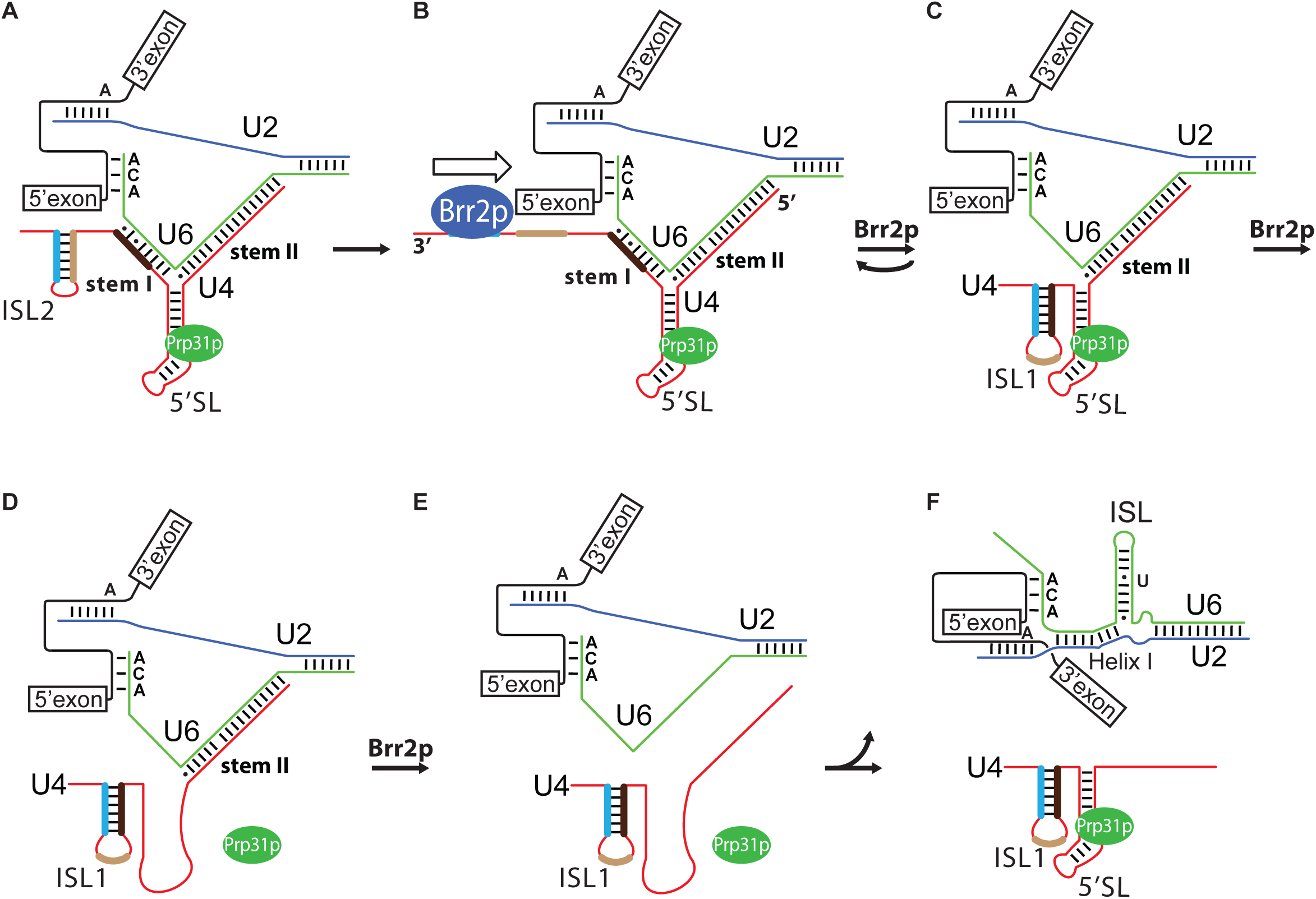
A model for the role of U4 in antagonizing and promoting Brr2p destabilization of U4/U6 stems I and II, as well as the intervening 5’SL. Panel A through panel F outlines a pathway for how Brr2p binds and likely pulls on U4 snRNA to unwind first stem I, which leads to formation of U4-ISL1 to prevent reannealing, and then further pulls to destabilize the U5-5’SL and stem II. (See Discussion for details.)

### Brr2p unwinds U4/U6 stem I in vivo

Brr2 unwinds the tri-snRNP and a minimal di-snRNP, and experiments with di-snRNAs implicated a role for Brr2p in acting on U4 with a 3’ to 5’ directionality and with direct unwinding of stem I (Mozaffari-Jovin et al. 2012; Raghunathan and Guthrie 1998; Theuser et al. 2016). Indeed, structures place Brr2p downstream of U4/U6 stem I, prior to U4/U6 unwinding (Plaschka et al. 2017; Zhan et al. 2018; Nguyen et al. 2016). In vivo, Brr2p cross-links to the U4 strand of stem I, consistent with a direct role in unwinding (Hahn et al. 2012). Here, we show that mutations that destabilize U4/U6 stem I suppress an unwinding-deficient *brr2* mutation, providing functional evidence in vivo that Brr2p unwinds U4/U6 stem I (Figure 1B).

### Evidence that reannealing of U4/U6 stem I antagonizes Brr2p-mediated unwinding

Unexpectedly, while some mutations in U4/U6 stem I suppressed the *brr2* mutation, as expected (Figure 1B), other mutations exacerbated the *brr2* mutation (Figure 2B), ultimately implicating a role for the mutually exclusive U4-ISL1 in productive U4/U6 unwinding. In a wild-type strain, disruption of ISL1 compromised growth as well as U4/U6 unwinding in isolated tri-snRNPs in vitro (Figure 3). Although a variation of ISL1 (hairpin II) had been proposed in free U4 snRNA (Myslinski and Branlant 1991)(Supplementary Figure 3), ISL1 had not yet been experimentally implicated in a spliceosome activation intermediate.

In principle, the U4-ISL1 could promote U4/U6 unwinding by directly activating Brr2p, but this seems unlikely, given Brr2p must already be active to unwind U4/U6 stem I, a prerequisite for U4-ISL1 formation at this stage. Formally, the ISL1 could also promote unwinding by somehow activating the U4 substrate, but again U4-ISL1 would form after U4/U6 stem I unwinding – as well as downstream of Brr2p, whereas the remainder of the U4 structures to be disrupted would be upstream of Brr2p. Further, the downstream strand of U4-ISL1 is unlikely to promote U4/U6 stem I unwinding directly by toehold-mediated strand displacement (Rodgers et al. 2016), given insufficient complementarity for a toehold and given Brr2p binding to this downstream strand before unwinding begins (Plaschka et al. 2017).

Alternatively, given that the formation of U4-ISL1 should liberate free energy, as previously suggested (Myslinski and Branlant 1991), and given the mutual exclusivity between U4-ISL1 and U4/U6 stem I, our data is consistent with a role for U4-ISL1 in driving the U4/U6 unwinding reaction forward by antagonizing the reverse reaction of U4/U6 stem I reannealing, which implies that unwinding of U4/U6 stem I is not inherently irreversible.

This model is particularly compelling, because it is otherwise difficult to rationalize why perturbation to a product of Brr2p-mediated U4/U6 stem I unwinding (i.e., U4-ISL1) could impact Brr2p-mediated unwinding. Further, our initial genetic results can be understood in terms of this model: i) specific destabilization by mutation of U4/U6 stem I favors the stability of the products state (U4-ISL1) and suppresses a *brr2* mutation, whereas specific destabilization of U4-ISL1 favors the substrate state (U4/U6 stem I) and exacerbates a *brr2* mutation (Figure 2A-D). Further genetic results strongly support such a role for the U4-ISL1 by providing evidence for the importance of balancing the stabilities of U4/U6 stem I and U4-ISL1. Specifically, the combination of U4-C81G or U4-C81A, which both specifically destabilize U4-ISL1 and exacerbate *brr2-R1107L*, with U4-U57G, which specifically destabilizes U4/U6 stem I and suppresses *brr2-R1107L*, result in an intermediate, wild-type phenotype, consistent with a rebalancing of the relative stabilities of the two structures and with equilibration between U4/U6 stem I and U4-ISL1 (Figure 2D).

Previous data implicated a role for the U6-ISL in driving U4/U6 stem II unwinding, just as we have implicated a role for U4-ISL1 in driving U4/U6 stem I unwinding. Specifically, a mutant in U4 that hyperstabilizes U4/U6 stem I was suppressed by a double mutation in U6 that hyperstabilizes U6-ISL, the product of unwinding U4/U6 stem II, suggesting that unwinding of U4/U6 stem II is reversible and could be promoted by U6-ISL (Kuhn et al. 2002). However, we show here that one of these mutations in U6 (A62U) suppresses a *brr2* mutation defective in U4/U6 unwinding by destabilizing U4/U6 stem I (Figure 1C), and in addition multiple tests revealed no evidence that weakening the U6 ISL compromised the *brr2* mutation (Figure Supplementary 5), arguing that the U6 ISL is not in equilibrium with U4/U6 during spliceosome activation, and that the U6 ISL does not drive unwinding.

Formation of U4-ISL1 might allow the U6 nucleotides no longer base paired with U4 to find U2 and form helix I during spliceosome activation, and biochemical data for the minor spliceosome in humans have demonstrated a spliceosomal complex in which U4atac/U6atac stem I is unwound and U12/U6atac helix Ia, the analog of U2/U6 helix Ia, is formed, while U4atac/U6atac stem II remains intact (Frilander and Steitz 2001). However, structural data on the major spliceosome during Bact formation (Townsend et al. 2020) provided evidence that U6-ISL is formed prior to U2/U6 helix I, suggesting that U4/U6 stem II needs to be unwound before U2/U6 helix I can form, consistent with our failure to find evidence that U2/U6 helix Ia formation helps to drive U4/U6 stem I unwinding (Figure Supplementary 5).

### Formation of U4-ISL2 may control Brr2p binding and antagonize U4-ISL1 during U4/U6 annealing

Genetic dissection of U4-ISL1 led us to the discovery of another intramolecular stem loop, U4-ISL2. Specifically, mutations that disrupt U4-ISL2 suppress a *brr2* mutation, in contrast to mutations that disrupt U4-ISL1, which antagonize the same *brr2* mutation (Figure 4C vs 2B). This genetic interaction suggests U4-ISL2 may inhibit recruitment of Brr2p to U4 until the stage of spliceosome activation, given sequestration of the Brr2p binding site in U4 by U4-ISL2, as proposed for U4/U6 stem III in humans (Charenton et al. 2019; Zhang et al. 2024; Jakab et al. 1997). Indeed, whereas U4-ISL2 is mutually exclusive with U4-ISL1, ISL2 can form when U4/U6 stem I is present, consistent with a role for ISL2 in preserving U4/U6 stem I (Figure 4A). Thus, U4-ISL2 may inhibit Brr2p and preserve U4/U6 stem I, both by preventing U4 binding by Brr2p and by antagonizing U4-ISL1 formation. However, after U4/U6 stem I unwinding, ISL1 likely dominates over U4-ISL2, because ISL1 is more stable than ISL2. Alternatively, or in addition, U4-ISL2 may promote reannealing of U4/U6 by antagonizing U4-ISL1 and thereby enabling formation of the mutually exclusive U4/U6 stem I.

### Brr2p unwinds both the U4-5’SL, releasing bound proteins, and U4-U6 stem II

Brr2p alone unwinds naked U4/U6 in vitro (Mozaffari-Jovin et al. 2012), implying translocation along U4 to the 5’ end and direct disruption of both stem I and stem II. However, in the context of a ribonucleoprotein complex, unwinding of U4/U6 may require a different mechanism. Indeed, the binding of Snu13, Prp31, and Prp3 to the intervening 5’SL, for example, inhibits Brr2-mediated unwinding of U4/U6 in vitro (Theuser et al. 2016). Further, in vivo cross-linking data implicates direct unwinding of U4/U6 stem I but not of stem II (Hahn et al. 2012). Nevertheless, our genetic results implicate Brr2p in unwinding of both stem I and stem II, raising questions as to how stem II is unwound.

Three models have been considered/proposed for the unwinding of U4/U6 stem II (Nielsen and Staley 2012; Theuser et al. 2016). First, Brr2p translocates processively through to the 5’ end of U4, directly disrupting stem I, the 5’SL, and stem II, as inferred in vitro with naked RNA. Second, Brr2p translocates through stem I, unwinding stem I directly, but because Brr2p is anchored in the context of a ribonucleoprotein particle, Brr2p pulls on U4, unwinding stem II indirectly; in this model Brr2p could unwind the 5’SL directly or indirectly by pulling. Third, Brr2p, after directly unwinding U4/U6 stem I, bypasses the 5’SL and then translocates along U4 again to disrupt U4/U6 stem II directly. The first two models predict Brr2p destabilizes the 5’SL and dissociates bound proteins (Snu13p and Prp31p); the third model does not. Our analysis of U4/U6 unwinding in the context of U4/U6-U5 snRNPs reveals the release of Prp31p from U4 snRNA, ruling out the third model and supporting the first and second models. Crosslinking of Brr2p to the 3’ side of the 5’SL could be consistent with either the first or the second model, with Brr2p unwinding directly or indirectly by pulling. However, in the spliceosomal B complex, in which Brr2 unwinds U4/U6, Brr2p is anchored by extensive protein-protein and RNA-protein interactions (Plaschka et al. 2017), and Brr2p does not crosslink to U4/U6 stem II in vivo (Hahn et al. 2012). These observations favor the second model over the first model, that Brr2p translocation along U4 leads to pulling of U4 and indirect unwinding of U4/U6 stem II.

### Brr2p unwinds U4/U6 stem I and stem II in a manner that is promoted by U4-ISL1

We envision a model for Brr2p mediated unwinding as described below (Figure 7). This model is in part based on the recent findings that isolation of budding yeast spliceosomal complexes at low ATP may be off-pathway complexes, a finding that has implications for several structures of yeast spliceosomes that have been interpreted as indicating an activation mechanism that is different from the activation mechanism of mammalian spliceosomes, with respect to when U6 binds a 5’ splice site and how Brr2p is upregulated for unwinding; indeed, these recent findings suggest that yeast and humans share an evolutionary common pathway of spliceosomal activation (Fu and Hoskins 2025). Our model for Brr2p mediated unwinding is therefore set in the context of mammalian spliceosomal activation. In a B-like spliceosomal complex, the starting point for the model (Figure 7A), Brr2p binding to U4 is antagonized by ISL2 (Figure 4), just as Brr2 binding is antagonized in humans by a U4/U6 duplex called stem III (Charenton et al. 2019); additionally, the U6 ACAGA sequence has bound to the 5’SS, as shown for humans (Zhang et al. 2024). Then, the U4-ISL2 unwinds, and Brr2p binds U4 downstream of U4/U6 stem I, after which Brr2p pulls U4, translocating along U4 in a 3’ to 5’ direction but moving U4, relative to the spliceosome in a 5’ to 3’ direction (Figure 7B). Brr2p pulling directly collides with and unwinds U4/U6 stem I, after which U4-ISL1 forms, thereby antagonizing reannealing of stem I (Figure 1-3; Figure 7C). Brr2p continues pulling, first destabilizing the U4-5’SL and liberating bound proteins (e.g., Prp31p, Figure 6) and then destabilizing stem II indirectly by continued pulling by Brr2p (Figure 5; Figure 7D, E).

Liberated U6 snRNA forms the U6-ISL and helix I with U2, while free U4 rebinds proteins at the U4-5’SL (i.e., Snu13p and Prp31p; Figure 7F). By this pathway, Brr2p unwinds both U4/U6 stems I and II in a manner promoted by the U4-ISL1, which prevents reannealing. Phylogenetic comparisons imply that this role for the U4-ISL1 is conserved from budding yeast to humans (Figure Supplementary 3).

## Materials and Methods

### Yeast manipulations

Yeast transformations were performed using the LiOAc method (Gietz and Schiestl 2007). After transformation, 5-fluoroorotic acid (5-FOA) was used to counter-select against yeast that had not passively lost the covering *URA3* plasmid. Yeast growth assays were performed by serial diluting, in 10-fold increments, yeast from overnight cultures grown in yeast peptone dextrose supplemented with adenine (YPDA; 1% yeast extract, 2% peptone, 2% dextrose, 0.004% adenine sulfate) or minimal media when indicated and then spotting with a frogger on solid YPDA or solid 5-FOA medium (5-FOA; 0.67% yeast nitrogen base, 0.2% drop-out mix—uracil, 2% glucose, 50 µg/mL uracil, 0.1% 5-FOA, and 2% agar) at the indicated temperatures.

### Yeast strains and plasmids

See table 1 for list of yeast strains and plasmids. The yeast strain yJPS1589, quadruply deleted for *SNR20, SNR14, SNR6, and BRR2,* was constructed as follows. The parental yeast strain yJPS628 encoded deletions for *snr20*::*LYS2, snr14*::*kanMX4* and *snr6*::*LEU2* and contains a *URA3* plasmid, pJPS467, expressing WT *SNR20*, *SNR14* and *SNR6*. The strategy was i) to regain the *LEU2* marker to allow use of the *BRR2* expressing plasmids (pJPS3086 and pJPS3087) and ii) to delete *BRR2* using a hygromycin B module (pJPS2231-hphMX4). In the first step of regaining the *LEU2* marker, yJPS628 was transformed with pJPS113 (*SNR20-HIS3*), pJPS464 (*SNR14-ADE2*) and pJPS12 (*SNR6-TRP1*), and yeast colonies that had passively lost the *URA3* plasmid, pJPS467, were selected. This yeast strain was subsequently transformed with a PCR product made using a hygromycin B module (pJPS2231-hphMX4) as template and oligonucleotide primers (5’U6-hph and 3’U6-hph) having *SNR6* overhangs of sequences upstream and downstream of *snr6*::*LEU2*; recombined transformants were selected on solid YPDA supplemented with 200 μg/ml hygromycin B. The resulting yeast strain was confirmed for loss of *LEU2* (*snr6*::*LEU2* replaced with *snr6*::hphMX4) by replica to CSM-leu and YPDA+ hygromycin B. Because the hphMX4 was needed for disruption of *BRR2* at a later step, we regained hphMX4 cassette by making use of the hisG-URA3-hisG approach (Alani et al. 1987). Yeast was transformed with a PCR product made using a hisG::*URA3*::hisG (pJPS644/pWY203) template and oligonucleotide primers (5’hph-hisG and 3’hph-hisG) having hphMX4 overhangs and selected for Ura+ Hyg-yeast. To regain the *URA3* marker the yeast was subsequently counter-selected on 5-FOA. In the second step of deleting *BRR2*, the resulting yeast strain was transformed first with a *BRR2-LEU2* plasmid (pJPS3086) and then with a PCR product made using a hygromycin B module (pJPS2231-hphMX4) as template and oligonucleotide primers (5’Brr2-hph and 3’Brr2-hph) having overhangs of sequences upstream or downstream of genomic *BRR2*; transformants were selected on YPD supplemented with 200 μg/ml hygromycin B. The final strain was transformed with pJPS3030 (*URA3, BRR2, U6, U4, U2*) and selected after growth in CSM-ura for passive loss of *BRR2-LEU2, SNR20-HIS3*, *SNR14-ADE2* and *SNR6-TRP1* by replica to CSM-ade-leu-his-trp and CSM-ura.

**Table 1.** Yeast strains and plasmids.

| Strain | Genotype | Reference |
| --- | --- | --- |
| yJPS628 | *MATa ade2-101; his3Δ200; leu2Δ1; lys2-801; trp1Δ63; ura3-52; snr6::LEU2; snr14::kanMX4; snr20::LYS2; [pJPS467]* | (Bordonne et al. 1990) |
| yJPS629 | *MATa his3Δ1; leu2Δ0; lys2Δ0; ura3Δ0; snr14::LYS2; [pYCp50-SNR14]* | (Bordonne et al. 1990) |
| yJPS776 | *MATa his3Δ1; leu2Δ0; met15Δ0; ura3Δ0; BRR2-TAP::kanMX6* | (Bellare et al. 2008) |
| yJPS788/PRY118 | *MATa brr2::LEU2; ade2; lys2; his3; ura3; leu2; [pJPS478-BRR2]* |  |
| yJPS1589 | *MATa ade2-101; his3Δ200; leu2Δ1; lys2-801; trp1Δ63; ura3-52; snr6::hisG; snr14::kanMX4; snr20::LYS2; brr2::hphMX4; [pJPS3030]* | this study |
| yJPS1588 | *MATa his3Δ1; leu2Δ0; lys2Δ0; ura3Δ0; snr14::LYS2; ade::hisG; brr2::kanMX6; [pJPS3031]* | this study |
| YGR091W | *MATa his3Δ1; leu2Δ0; met15Δ0; ura3Δ0; PRP31::TAP-HIS3MX6* | (Ghaemmaghami et al. 2003) |
| Plasmid | Description | Reference |
| pYCp50-SNR14 | URA3 plasmid expressing SNR14 | (Bordonne et al. 1990) |
| pJPS12 | TRP1 plasmid expressing SNR6 | (Madhani et al. 1990) |
| pJPS113 | HIS3 plasmid expressing SNR20 without the fungal domain | (Shuster and Guthrie 1990) |
| pJPS434/pyU4wt | HIS3 plasmid expressing SNR14 | (Bordonne et al. 1990) |
| pJPS464 | ADE2 plasmid expressing SNR14 from pASZ11(Stotz and Linder 1990) | (Hilliker and Staley 2004) |
| pJPS467 | *URA3* plasmid expressing *SNR20, SNR14, SNR6* | (Hilliker and Staley 2004) |
| pJPS478/pPR130 | *HIS* plasmid expressing *BRR2* | (Raghunathan and Guthrie 1998) |
| pJPS619/pSN108 | *URA3* plasmid expressing *BRR2* | (Noble and Guthrie 1996) |
| pJPS644/pWY203 | Integrating plasmid that contains *hisG-URA3-hisG* | (Galli and Schiestl 1998) |
| pJPS2433 | *HIS3* plasmid expressing *brr2-R1107L* | (Zhao et al. 2009) |
| pJPS2231-hphMX4 | Hygromycin cassette | (Tagwerker et al. 2006) |
| pJPS2579 | *HIS* plasmid expressing TAP-tagged *BRR2* | this study |
| pJPS3030 | *URA3* plasmid expressing *SNR20, SNR14, SNR6, BRR2* | this study |
| pJPS3031 | *URA3* plasmid expressing *BRR2* and *SNR14* | this study |
| pJPS3032 | *HIS3* plasmid expressing *SNR14* | this study |
| pJPS3086 | *LEU2* plasmid expressing *BRR2* | this study |
| pJPS3087 | *LEU2* plasmid expressing *brr2-R1107L* | this study |
| pJPS3088 | *ADE2* plasmid expressing U4-U57G | this study |
| pJPS3089 | *ADE2* plasmid expressing U4-U57C | this study |
| pJPS3090 | *ADE2* plasmid expressing U4-U57A | this study |
| pJPS473 | *ADE2* plasmid expressing U4-G58C | Gift of AK.Hilliker |
| pJPS469 | *ADE2* plasmid expressing U4-G58U | Gift of AK.Hilliker |
| pJPS468 | *ADE2* plasmid expressing U4-G58A | Gift of AK.Hilliker |
| pJPS3008 | *TRP1* plasmid expressing U6-A62U | this study |
| pJPS3009 | *TRP1* plasmid expressing U6-A62C | this study |
| pJPS474 | *ADE2* plasmid expressing U4-C59A | (Hilliker and Staley 2004) |
| pJPS475 | *ADE2* plasmid expressing U4-U60C | (Hilliker and Staley 2004) |
| pJPS1992 | *ADE2* plasmid expressing U4-G61A | Gift of MA.Mefford |
| pJPS1974 | *ADE2* plasmid expressing U4-G62C | Gift of MA.Mefford |
| pJPS1975 | *ADE2* plasmid expressing U4-U63A | Gift of MA.Mefford |
| pJPS1996 | *ADE2* plasmid expressing U4-U64A | Gift of MA.Mefford |
| pJPS1997 | *ADE2* plasmid expressing U4-U64G | Gift of MA.Mefford |
| pJPS3075 | *ADE2* plasmid expressing U4-U64C | this study |
| pJPS3091 | *ADE2* plasmid expressing U4-C82G | this study |
| pJPS3092 | *ADE2* plasmid expressing U4-C82A | this study |
| pJPS3093 | *ADE2* plasmid expressing U4-C82U | this study |
| pJPS3094 | *ADE2* plasmid expressing U4-G61A C82G | this study |
| pJPS3095 | *ADE2* plasmid expressing U4-G61A C82A | this study |
| pJPS3096 | *ADE2* plasmid expressing U4-G61A C82U | this study |
| pJPS3097 | *ADE2* plasmid expressing U4-G61U C82G | this study |
| pJPS3098 | *ADE2* plasmid expressing U4-G61U C82A | this study |
| pJPS3099 | *ADE2* plasmid expressing U4-G61U C82U | this study |
| pJPS3100 | *ADE2* plasmid expressing U4-G61C C82G | this study |
| pJPS3101 | *ADE2* plasmid expressing U4-G61C C82A | this study |
| pJPS3102 | *ADE2* plasmid expressing U4-G61C C82U | this study |
| pJPS2001 | *ADE2* plasmid expressing U4-G62U | this study |
| pJPS3103 | *ADE2* plasmid expressing U4-G62U C81A | this study |
| pJPS3104 | *ADE2* plasmid expressing U4-C81A | this study |
| pJPS3105 | *ADE2* plasmid expressing U4-CCA81AAC | this study |
| pJPS3106 | *ADE2* plasmid expressing U4-79uuaac | this study |
| pJPS3107 | *ADE2* plasmid expressing U4-79uuaaca | this study |
| pJPS3108 | *ADE2* plasmid expressing U4-69aaa | this study |
| pJPS3109 | *ADE2* plasmid expressing U4-74aa | this study |
| pJPS3110 | *ADE2* plasmid expressing U4-69aaa74aa | this study |
| pJPS3111 | *ADE2* plasmid expressing U4-72gg | this study |
| pJPS3112 | *ADE2* plasmid expressing U4-72cc | this study |
| pJPS2851 | *ADE2* plasmid expressing U4-U2G | this study |
| pJPS2767 | *ADE2* plasmid expressing U4-C3U | this study |
| pJPS3113 | *ADE2* plasmid expressing U4-C4A | this study |
| pJPS2826 | *ADE2* plasmid expressing U4-U5G | this study |
| pJPS3114 | *ADE2* plasmid expressing U4-U5C | this study |
| pJPS3115 | *ADE2* plasmid expressing U4-U6A | this study |
| pJPS3116 | *ADE2* plasmid expressing U4-U8C | this study |
| pJPS3117 | *ADE2* plasmid expressing U4-A11G | this study |
| pJPS2858 | *ADE2* plasmid expressing U4-A11U | this study |
| pJPS3118 | *ADE2* plasmid expressing U4-C12A | this study |
| pJPS2765 | *TRP1* plasmid expressing U6-A75U | this study |
| pJPS2766 | *TRP1* plasmid expressing U6-A75C | this study |
| pJPS2865 | *TRP1* plasmid expressing U6-A75G | this study |
| pJPS2763 | *TRP1* plasmid expressing U6-A73G | this study |
| pJPS2764 | *TRP1* plasmid expressing U6-A73C | this study |
| pJPS2864 | *TRP1* plasmid expressing U6-A73U | this study |
| pJPS3055 | *ADE2* plasmid expressing U4-GG61CC | this study |
| pJPS3051 | *ADE2* plasmid expressing U4-CC81GG | this study |
| pJPS3057 | *ADE2* plasmid expressing U4-GG61CC CC81GG | this study |
| pJPS3119 | *ADE2* plasmid expressing U4-A20G | this study |
| pJPS3120 | *ADE2* plasmid expressing U4-U53C | this study |
| pJPS3121 | *ADE2* plasmid expressing U4-20G 53C | this study |
| pJPS3122 | *ADE2* plasmid expressing U4-AC20CG | this study |
| pJPS3123 | *ADE2* plasmid expressing U4-GU52CG | this study |
| pJPS3124 | *ADE2* plasmid expressing U4-20CG 52CG | this study |
| pJPS3125 | *ADE2* plasmid expressing U4-20CG 52CA | this study |
| pJPS3126 | *ADE2* plasmid expressing U4-20GA 52CG | this study |
| pJPS3127 | *ADE2* plasmid expressing U4-20GA | this study |
| pJPS3128 | *ADE2* plasmid expressing U4-52CA | this study |
| pJPS3129 | *ADE2* plasmid expressing U4-U60C A83G | this study |
| pJPS3130 | *ADE2* plasmid expressing U4-U60C A83C | this study |
| pJPS3131 | *ADE2* plasmid expressing U4-U60C A83U | this study |
| pJPS3132 | *ADE2* plasmid expressing U4-A83G | this study |
| pJPS3133 | *ADE2* plasmid expressing U4-A83C | this study |
| pJPS3134 | *ADE2* plasmid expressing U4-A83U | this study |
| pJPS3000 | *ADE2* plasmid expressing U4-G62C C81U | this study |
| pJPS2959 | *ADE2* plasmid expressing U4-C81U | this study |
| pJPS3135 | *ADE2* plasmid expressing U4-C81G | this study |
| pJPS3136 | *ADE2* plasmid expressing U4-U57G C81G | this study |
| pJPS3137 | *ADE2* plasmid expressing U4-C81G G62C | this study |
| pJPS3138 | *ADE2* plasmid expressing U4-C81G G62U | this study |
| pJPS3139 | *ADE2* plasmid expressing U4-C81A G62C | this study |
| pJPS3140 | *ADE2* plasmid expressing U4-C81A G62U | this study |
| pJPS2004 | *ADE2* plasmid expressing U4-U63G | Gift of MA.Mefford |
| pJPS3141 | *ADE2* plasmid expressing U4-A80U | this study |
| pJPS3142 | *ADE2* plasmid expressing U4-A80C | this study |
| pJPS3143 | *ADE2* plasmid expressing U4-A80U U63A | this study |
| pJPS3144 | *ADE2* plasmid expressing U4-A80U U63G | this study |
| pJPS3145 | *ADE2* plasmid expressing U4-A80C U63A | this study |
| pJPS3146 | *ADE2* plasmid expressing U4-A80C U63G | this study |
| pJPS2827 | *ADE2* plasmid expressing U4-C12U | this study |
| pJPS3147 | *ADE2* plasmid expressing U4-U57G C81A | this study |
| pJPS3148 | *ADE2* plasmid expressing U4-C12G | this study |
| pJPS1985 | *TRP1* plasmid expressing U6-C33G G55C | Gift of MA.Mefford |
| pJPS3149 | *TRP1* plasmid expressing U6-A82U | this study |
| pJPS3150 | *TRP1* plasmid expressing U6-A82C | this study |
| pJPS3151 | *TRP1* plasmid expressing U6-C84A | this study |
| pJPS3152 | *TRP1* plasmid expressing U6-C84G | this study |
| pJPS3153 | *TRP1* plasmid expressing U6-C84U | this study |
| pJPS2860 | *TRP1* plasmid expressing U6-C67U | this study |
| pJPS2861 | *TRP1* plasmid expressing U6-C69U | this study |
| pJPS2862 | *TRP1* plasmid expressing U6-C69A | this study |
| pJPS2863 | *TRP1* plasmid expressing U6-C69G | this study |
| pJPS410 | *HIS3* plasmid expressing U2-C14G C15G | Gift of AK.Hilliker |
| pJPS3154 | *HIS3* plasmid expressing U2-C15G | this study |

The yeast strain yJPS1588, doubly deleted for *BRR2* and *SNR14*, was constructed as follows. The strategy was to i) delete *ADE2* using the hisG-URA3-hisG approach (Alani et al. 1987), so a library of *U4-ADE2* plasmids could be used in the strain and ii) delete *BRR2* using the kanMX4 cassette. The parental yeast strain yJPS629 is deleted for *snr14*::*LYS2* and contains a *URA3* plasmid, pYCp50-*SNR14* (Bordonne et al. 1990), expressing *SNR14* and was generated in the Guthrie lab from the Research Genetic Consortium yeast strain BY4743, which was generated from a cross between BY4741 and BY4742 (Brachmann et al. 1998). In the first step of deleting *ADE2*, we needed to start with rendering the *URA3* marker available. Strain yJPS629 was transformed with a WT *SNR14*-*HIS3* plasmid (pJPS3032) and counter-selected on 5-FOA to isolate strains that lost the *URA3* plasmid. We then used a split-marker approach (Hoogt et al. 2000) to obtain PCR products for hisG-URA3-hisG with sequences upstream or downstream of *ADE2*. The yeast strain was transformed with two PCR products generated using hisG::*URA3*::hisG (pJPS644-pWY203) as template and two sets of primers (primer set 1: 5’ade2-hisG and URA3 R; primer set 2:

URA3 F and 3’ade2-hisG); transformants were selected on -ura medium. PCR- and auxotrophic-verified ade-yeast were counter-selected on 5-FOA to regain the *URA3* marker by selection for hisG recombinants. In the second step of deleting *BRR2*, the yeast strain was transformed with pJPS3031 (*SNR14-BRR2-URA3*) and subsequently transformed with a PCR product generated using genomic DNA isolated from a *BRR2* heterozygote from the Yeast Deletion Collection (Giaever and Nislow 2014) (yJPS635/BY4743 *brr2*::*kanMX4*) as template with oligonucleotide primers (5’BRR2-kan and 3’BRR2-kan) containing sequences upstream or downstream of *BRR2* and subsequently selected on YPD supplemented with 500 μg/ml G418.

#### pJPS2579

The TAP-kanMX fragment was obtained by PCR on genomic DNA from yJPS776; this fragment was then transformed into yJPS788, expressing the only copy of *BRR2* from pJPS478. The resulting strain was selected for growth on YPD supplemented with 500 μg/ml G418. The plasmid was recovered and the 3’ end of *BRR2* containing the TAP-kanMX integrated fragment was sequenced to confirm that the TAP sequence was in frame.

#### pJPS3030

The covering plasmid expression *BRR2*, *SNR14*, *SNR6* and *SNR20* genes was made by digesting pJPS619 with SmaI, heat inactivating SmaI, CIP treating, and then ligating to a blunt PCR product amplified using pJPS467, encoding *SNR14, SNR6* and *SRN20*, as template and oligos 5’U4U6U2 and 3’U4U6U2 as primers.

#### pJPS3031

The covering plasmid with both *BRR2* and *SNR14* was made by first PCR amplifying *SNR14* using pJPS434 (pyU4wt (Bordonne et al. 1990)) as template and oligonucleotide primers (5’U4 and 3’U4), which both contain an XmaI digested site. Then, this PCR product and pJPS619 were cut with XmaI and ligated.

#### pJPS3032

pRS313 (Sikorski and Hieter 1989) (*HIS3*) vector plasmid and pJPS434 (pyU4wt (Bordonne et al. 1990)), expressing *SNR14*, were both digested with EcoRI and BamHI and ligated.

#### pJPS3086

pRS315 (Sikorski and Hieter 1989) (*LEU2*) vector plasmid was digested with SacI, which was followed by heat inactivating SacI, CIP treatment, and then ligation to a SacI *BRR2* fragment from pPR130 (Raghunathan and Guthrie 1998).

#### pJPS3087

pRS315 (Sikorski and Hieter 1989) (*LEU2*) vector plasmid was digested with SacI, which was followed by heat inactivating SacI, CIP treatment, and then ligation to a SacI *brr2-R1107L* fragment from pJPS2433.

The pJPS464 plasmid that expresses wild-type *SNR14* from pASZ11 was used as the parental plasmid for all site-directed mutagenesis to generate the described *SNR14* mutations. The entire gene of *SNR14* was sequenced to verify that only the expected mutations were present.

Oligo list:

The underlined sequences indicate restriction site sequences.

5’U6-hph

5’GCCTGGCATGAACAGTGGTAAAAGTATTTCGTCCACTATTTTCGGCTACTATAAATAAATGCGTACGCTGCAGGT CGAC

3’U6-hph

5’CTATTCGAGATGTTAGTTATCTATATCATACAGGAAGATAAAGATACACTGCTGTACGCTGGATCGATGAATTCG

AGCTCG

5’Brr2-hph

5’GAAGGTAAATATAAGTAATTGCTTTGGAAGATTTACCGTGAGCCTTCGTTATTAAAACGTACGCTGCAGGTCGAC

3’Brr2-hph

5’CGCAAGAGAATGTTATATATTGAAATCCATTCGATTATCCAGGACTAAACAATGATTATCGATGAATTCGAGCTCG

5’hph-hisG

5’ACAGGGGTTCACATCATCACCACCATCATGACTACGATATACCCACAACCGCCAGCGAGCTATTGAGGCCAATTTTTCCTTTGACGAGC

3’hph-hisG

5’TATCGAATCGACAGCAGTATAGCGACCAGCATTCACATACGATTGACGCATGATATTACGGCTAAGATAATGGGGCTCTTTACATTTCC

5’U4U6U2 CCTGCAGGTCGACTCTAGAG 3’U4U6U2 CATTCGTAAGGGCCAATTC 5’U4

GAAACCCGGGGATATCATAAGTGCGTTCGAGG

3’U4

GAAACCCGGGAATTCGGTGAAAAAGAAAAGAAAAATATGG

5’ade2-hisG

TTCTAAGTACATCCTACTATAACAATCAAGAAAAACAAGAAAATCGGACAAAACAATCAAGTATGGCAAAGTTGCCAAGATCTTCCAGTG

URA3 R TAGCAGCACGCCATAGTGACTG

3’ade2-hisG

GAACAGCGGATCCTCAAAAATTCGTAATATATAAGTTTATTGATATACTTGTACAGCAAATAATTATAAAATGATATACCTATTTTTTAG

URA3 F GGAGCACAGACTTAGATTGGTATATATACGC 5’BRR2-kan

GCGTGTTTGCTTCTTTGACGATC

3’BRR2-kan GGAGAACAGGATGAAGATGAAGATG

### Isolation and unwinding of tri-snRNPs

The yeast strains expressing either TAP tagged Brr2p (yJPS776) or TAP tagged Prp31p (YGR091W) were used for isolating tri-snRNPs having Brr2p or Prp31p TAP tagged, respectively. To isolate tri-snRNPs containing different U4 mutations, yJPS1588 was transformed with pJPS2579 (*HIS3-BRR2-TAP*) and pJPS464 (*ADE2-SNR14*) or *snr14* with the indicated U4 mutation and selected for passive loss of the *URA3* plasmid by replica plating to CSM-ade-his and CSM-ura. The strains were grown in YPDA to late-log phase at 30 °C (OD600 ∼ 2.0), and cells were harvested by spinning at 4.5K for 15’ in a Sorvall RC-3B (Small et al. 2006). Cell pellets were washed with ice-cold 100 ml of AGK buffer (10 mM HEPES pH 7.9, 1.5 mM MgCl_2_, 200 mM KCl, 10% vol/vol glycerol, 0.5 mM DTT) per liter of yeast culture and collected by spinning 4.5K for 15’ in an RC-3B. The cells were resuspended in 20 ml ice-cold AGK buffer and transferred to a 50 ml canonical tube. The cells were collected by spinning 2.5K for 10’ in an RC-3B and resuspended in 0.8 volume of ice-cold AGK buffer containing EDTA-free protease inhibitors (Roche). The yeast suspension was added dropwise into a 1 L glass beaker with ∼200 ml liquid nitrogen and the pellets were transferred to a 50 ml canonical and stored at −80 °C or immediately processed, for cell lysis, using a liquid nitrogen cooled freezer-mill (6875 Spex SamplePrep, Cole-Parmer) with the following parameters: 5 cycles of 3 min with 15 cycles per second and 1 min pause in between. The powder was thawed at 4 °C and centrifugated at 17K for 30 minutes in an SS34 rotor at 4 °C. The supernatant was carefully transferred to 70.1 Ti tubes and the lysate was clarified by spinning at 37K for 1.25 hr at 4 °C. The clear fraction was collected, carefully avoiding the bottom pellet and the top lipid layer. Then, 1 ml of extract was mixed with 200 µl IgG beads (Cytiva) that had been washed 4x with 1ml IPP150 buffer (10 mM Tris-HCl pH 8, 150mM NaCl, 0.1% NP-40, 10% glycerol) and incubated for 2 hrs with rotation at 4 °C. The IgG beads were washed 4x with 1.5 ml buffer D (20 mM Hepes pH 7.9, 0.2 mM EDTA, 50 mM KCl, 0.5 mM DTT, 20% glycerol), resuspended in 160 µl buffer D and 40 µl aliquots were flash frozen in liquid nitrogen and stored at −80°C.

Unwinding of isolated tri-snRNPs was done at 15 °C under splicing conditions (3% PEG, 60 mM KH_2_PO_4_/K_2_HPO_4_ pH 7.0, 2.5 mM MgCl_2_, and 40% buffer D derived from the isolated tri-snRNP) and initiated with addition of 2 mM ATP. Aliquots of 20 µl were taken at indicated times and quenched by adding to a mix of cold 500 µl RNA buffer (50 mM Tris-HCl pH 7.4, 100 mM NaCl, 10 mM EDTA) and 500 µl cold acidic phenol:chloroform:isoamyl alcohol (125:24:1) at pH 4.5; cold solutions were used to maintain U4/U6 base pairing. RNA was extracted twice and precipitated with 1 ml EtOH containing 5 µl glycogen (10 mg/ml). The RNA was analyzed as described under northern blotting or primer extension.

Unwinding of isolated tri-snRNPs were performed to separate snRNA either remaining associated with Brr2p-TAP or Prp31p-TAP IgG beads (beads) or released (supernatant). The reactions contained 500 nM U4-5’SL as competitor were indicated; this RNA was heated to 70 °C for 5 minutes and then cooled on ice before adding to the reactions. The reactions in a volume of 30 µl were performed at 15 °C for 3 minutes under splicing conditions (3% PEG, 60 mM KH_2_PO_4_/K_2_HPO_4_ pH 7.0, 2.5 mM MgCl_2_, and 40% buffer D derived from the isolated tri-snRNP) and initiated with addition of 2 mM ATP final concentration. The reactions were pelleted by centrifugation for 15 seconds at 0.5 x g (Eppendorf centrifuge) and supernatant was quenched by adding to a mix of 300 µl RNA buffer (50 mM Tris-HCl pH 7.4, 100 mM NaCl, 10 mM EDTA) and 500 µl acidic phenol:chloroform:isoamyl alcohol (125:24:1) at pH 4.5. The beads were washed twice with 100 µl washing buffer (10 mM Tris-HCl pH 8.0, 100 mM NaCl, 2 mM MgCl_2_) that was added to the Eppendorf tube containing the quenched supernatant above. The IgG beads were quenched by adding to a mix of 500 µl RNA buffer (50 mM Tris-HCl pH 7.4, 100 mM NaCl, 10 mM EDTA) and 500 µl acidic phenol:chloroform:isoamyl alcohol (125:24:1) at pH 4.5. The RNA was isolated as described under primer extension.

#### Primer extension analysis

RNA was isolated from tri-snRNPs preparations, or total RNA was isolated from cells grown to log phase OD_600_=0.7-1.0 at 30 °C. RNA was isolated by hot acidic phenol:chloroform:isoamyl alcohol (125:24:1) at pH 4.5, and primer extensions were performed as described (Stevens and Abelson 2002) using Cy5-labeled oligos complementary to the five snRNAs. For experiments where U4-CC81GG was investigated all primer extension reactions were carried out using the U4-81 oligo instead of the ssU4 oligo, since the final two nucleotides in the ssU4 oligo no longer anneal to the mutated U4 sequence. Products were separated on a 6% denaturing polyacrylamide gel and scanned on a Typhoon (GE healthcare) and quantitated by Image J (NIH).

#### Primer list for primer extension

U1: snR19: ssU1 5’ GAATGGAAACGTCAGCAAACAC 3’

U2: snR20: ssU2M5: 5’ AAAGTCTCTTCCCGTCCATTTTATTA 3

U4: snR14: ssU4: 5’ ACCATGAGGAGACGGTCTGG 3’

U4: snR14: U4-81: 5’ TGACCATGAGGAGACGGTCT 3’

U5: snR7: ssU5-125: 5’ GGAGACAACACCCGGATGGTTCTGGTA 3’

U6: snR6: U63’: 5’ AACGAAATAAATCTCTTTGTAAAAC 3’

#### RT-PCR

Yeast was grown to early log phase at 30 °C and shifted to 16 °C for 3 hrs. RNA was prepared as for primer extension with an additional DNase treatment (Ambion) step. A total of 2 µg total RNA and 1 µg RT oligo 5’ TCGTACGCTTAGTTCATCCCA 3’ (*YOS1*) was heated to 70 °C for 5 minutes and put on ice. RT reaction was done using AMV RT according to manufacturer’s manual (Promega) and the PCR reaction was done using 1/3 volume cDNA and 2/3 volume PCR reaction (Platinum™ II Hot-Start PCR Master Mixes (2X)) according to manufacturer’s manual (Invitrogen) using a forward oligo 5’ AGGGGATATGGTACTGTTTGGA 3’ and the RT oligo for 18 cycles. Identical control reactions were carried out without AMV RT; these PCR reactions did not generate bands. Reactions were separated on a 2% agarose gel containing ethidium bromide (0.2 ng/ml) and bands were quantitated using Image J (NIH).

#### Northern blotting

Total RNA was extracted from 25–50 mL yeast cultures grown to an OD600 ∼ 0.7 using cold acidic phenol:chloroform:isoamyl alcohol (125:24:1) at pH 4.5; cold solutions were used to maintain U4/U6 base pairing. Then, 10 µg total RNA or RNA extracted from unwinding experiments (see above) was separated on a 6% native PAGE gel in 1x TBE and transferred semi-dry to Hybond-NX (Amersham) in 0.5x TBE. The membranes were cross-linked in a UV Stratalinker 2400 (Stratagene) using the auto crosslink feature. Membranes were blocked using 10 ml ExpressHyb Hybridization Solution (Takara) for 30 min after which 10 µl of 10 µM Cy5-U4 labeled probes (U4-14B: 5’ AGGTATTCCAAAAATTCCCTA 3’ and ssU4: 5’ ACCATGAGGAGACGGTCTGG 3’) was added for 1 hr in a hybridization oven set to 37 °C. The membranes were washed at room temperature four times with 2x SSC (20x SSC is 3 M NaCl, 0.3 M sodium citrate) and 0.1% SDS for 5 min and four times with 1x SSC and 0.1% SDS for 10 min. Membranes were scanned using a Typhoon (GE healthcare) and quantitated by Image J (NIH).

#### Conservation and alignment analysis

Alignment of the 177 seed sequences of U4 from RFam (Ontiveros-Palacios et al. 2024) (only 176 sequences were used since the U18778.1/15676-15446 sequence is identical to the AAEG01000006.1/103924-103694 sequence) was made using Clustal Omega (Madeira et al. 2024). The phylogenetic tree was visualized using iTol (Letunic and Bork 2024) and identified a node containing the related fungi species (*N. castellii*; AACF01000119.1/10356-10566*, S. pastorianus*; AF270843.1/940-1107*, S. kudriavzevii*; AACI02000576.1/2356-2546*, S. mikatae*; AABZ01000001.1/29367-29202/1-166*, S. paradoxus*; AABY01000063.1/21244-21029/1-216, and *S. cerevisiae*; URS00001143F5_4932/1-160) that was used in Supplemental Figure 3 together with selected species for comparison (*H. sapiens*; URS00003F07BD_9606/1-144*, P. troglodytes*; AC193264.3/174224-174084/1-141*, B. taurus*; AAFC03029538.1/15458-15318/1-141*, S. purpuratus*; AAGJ04107959.1/2135-2275*, G.aculeatus*; AANH01001405.1/82396-82256/1-141*, D. rerio*; BC124577.1/1-141*, D. melanogaster*; K03095.1/1-139*, S. pombe*; CU329671.1/467615-467481/1-135, and *V. Polyspora*; AAZN01000370.1/73687-73536/1-152). The node containing related fungi species contained two additional *S. cerevisiae* entries; U18778.1/15676-15446 and URS0000D77922_4932/1-161 that are not shown. The evolutionary distance (Jukes-Cantor) was calculated using the following website: https://www.nccbiology.com/support/242/.

## Supporting information

Nielsen-Suppl-all

## Acknowledgments

We thank all members of the Staley lab for critical comments to the manuscript. We thank Angela K. Hilliker and Melissa A. Mefford for plasmids. This work was supported by a National Institutes of Health (NIH) Research Project Grant (R01GM062264) awarded to J.P.S.

## References

Alani E, Cao L, Kleckner N. 1987. A Method for Gene Disruption That Allows Repeated Use of URA3 Selection in the Construction of Multiply Disrupted Yeast Strains. Genetics 116: 541–545. doi:10.1534/genetics.112.541

Bai R, Wan R, Yan C, Lei J, Shi Y. 2018. Structures of the fully assembled Saccharomyces cerevisiae spliceosome before activation. Science 360: 1423–1429. doi:10.1126/science.aau0325

Bellare P, Small EC, Huang X, Wohlschlegel J a, Staley JP, Sontheimer EJ. 2008. A role for ubiquitin in the spliceosome assembly pathway. Nat Struct Mol Biol 15: 444–51. doi:10.1038/nsmb.1401

Bertram K, Agafonov DE, Liu WT, Dybkov O, Will CL, Hartmuth K, Urlaub H, Kastner B, Stark H, Lührmann R. 2017. Cryo-EM structure of a human spliceosome activated for step 2 of splicing. Nature 542: 318–323. doi:10.1038/nature21079

Bordonne R, Banroques J, Abelson J, Guthrie C. 1990. Domains of yeast U4 spliceosomal RNA required for PRP4 protein binding, snRNP-snRNP interactions, and pre-mRNA splicing in vivo. Genes Dev 4: 1185–1196. doi:10.1101/gad.4.7.1185

Brachmann CB, Davies A, Cost GJ, Caputo E, Li J, Hieter P, Boeke JD. 1998. Designer deletion strains derived from Saccharomyces cerevisiae S288C: A useful set of strains and plasmids for PCR-mediated gene disruption and other applications. Yeast 14: 115–132. doi:10.1002/(SICI)1097-0061(19980130)14:2<115::AID-YEA204>3.0.CO;2-2

Brow DA. 2019. An Allosteric Network for Spliceosome Activation Revealed by High-Throughput Suppressor Analysis in Saccharomyces cerevisiae. Genetics 212: 111–124. doi:10.1534/genetics.119.301922

Brow DA, Guthrie C. 1988. Spliceosomal RNA U6 is remarkably conserved from yeast to mammals. Nature 334: 213–8. doi:10.1038/334213a0

Charenton C, Wilkinson ME, Nagai K. 2019. Mechanism of 5 ′ splice site transfer for human spliceosome activation. Science 364: 362–367. doi:10.1126/science.aax3289

Cordin O, Beggs JD. 2013. RNA helicases in splicing. RNA Biol 10: 83–95. doi:10.4161/rna.22547

Didychuk AL, Butcher SE, Brow DA. 2018. The life of U6 small nuclear RNA, from cradle to grave. RNA 24: 437–460. doi:10.1261/rna.065136.117

Fairman-Williams ME, Guenther UP, Jankowsky E. 2010. SF1 and SF2 helicases: Family matters. Curr Opin Struct Biol 20: 313–324. doi:10.1016/j.sbi.2010.03.011

Fica SM, Mefford MA, Piccirilli JA, Staley JP. 2014. Evidence for a group II intron-like catalytic triplex in the spliceosome. Nat Struct Mol Biol 21: 464–71. doi:10.1038/nsmb.2815

Fica SM, Oubridge C, Galej WP, Wilkinson ME, Bai X-C, Newman AJ, Nagai K. 2017. Structure of a spliceosome remodelled for exon ligation. Nature 542: 3–7. doi:10.1038/nature21078

Fica SM, Tuttle N, Novak T, Li N-S, Lu J, Koodathingal P, Dai Q, Staley JP, Piccirilli JA. 2013. RNA catalyses nuclear pre-mRNA splicing. Nature 503: 229–234. doi:10.1038/nature12734

Fortner DM, Troy RG, Brow DA. 1994. A stem/loop in U6 RNA defines a conformational switch required for pre-mRNA splicing. Genes Dev 8: 221–233. doi:10.1101/gad.8.2.221

Frilander MJ, Steitz JA. 2001. Dynamic exchanges of RNA interactions leading to catalytic core formation in the U12-dependent spliceosome. Mol Cell 7: 217–226. doi:10.1016/s1097-2765(01)00169-1

Fu X, Hoskins AA. 2025. Dynamics and evolutionary conservation of B complex protein recruitment during spliceosome activation. Nucleic Acids Res 53. doi:10.1093/nar/gkaf124.

Galej WP, Wilkinson ME, Fica SM, Oubridge C, Newman AJ, Nagai K. 2016. Cryo-EM structure of the spliceosome immediately after branching. Nature 537: 197–201. doi:10.1038/nature19316

Galli A, Schiestl RH. 1998. Effects of DNA Double-Strand and Single-Strand Breaks on Intrachromosomal Recombination Events in Cell-Cycle-Arrested Yeast Cells. Genetics 149: 1235–1250. doi:10.1093/genetics/149.3.1235

Ghaemmaghami S, Huh WK, Bower K, Howson RW, Belle A, Dephoure N, O’Shea EK, Weissman JS. 2003. Global analysis of protein expression in yeast. Nature 425: 737–741. doi:10.1038/nature02046

Giaever G, Nislow C. 2014. The yeast deletion collection: A decade of functional genomics. Genetics 197: 451–465. doi:10.1534/genetics.114.161620

Gietz RD, Schiestl RH. 2007. Quick and easy yeast transformation using the LiAc/SS carrier DNA/PEG method. Nat Protoc 2: 35–37. doi:10.1038/nprot.2007.14

Hahn D, Kudla G, Tollervey D, Beggs JD. 2012. Brr2p-mediated conformational rearrangements in the spliceosome during activation and substrate repositioning. Genes Dev 26: 2408–2421. doi:10.1101/gad.199307.112

Hang J, Wan R, Yan C, Shi Y. 2015. Structural basis of pre-mRNA splicing. Science 349: 1191–1198. doi:10.1126/science.aac8159

Hardin JW, Warnasooriya C, Kondo Y, Nagai K, Rueda D. 2015. Assembly and dynamics of the U4/U6 di-snRNP by single-molecule FRET. Nucleic Acids Res 43: 10963–10974. doi:10.1093/nar/gkv1011

Hilliker AK, Staley JP. 2004. Multiple functions for the invariant AGC triad of U6 snRNA. RNA 10: 921– 928. doi:10.1261/rna.7310704

de Hoogt R, Luyten WHML, Contreras R, De Backer MD. 2000. PCR- and Ligation-Mediated Synthesis of Split-Marker Cassettes with Long Flanking Homology Regions for Gene Disruption in Candida albicans. Biotechniques 28: 1112–16. doi:10.2144/00286st01

Hu JIM, Xu D, Schappert K, Xu YAN, Friesen JD. 1995. Mutational Analysis of Saccharomyces cerevisiae U4 Small Nuclear RNA Identifies Functionally Important Domains. Mol Cell Biol 15: 1274–1285. doi:10.1128/MCB.15.3.1274

Jakab G, Mougin A, Kis M, Pollák T, Antal M, Branlant C, Solymosy F. 1997. Chlamydomonas U2, U4 and U6 snRNAs. An evolutionary conserved putative third interaction between U4 and U6 snRNAs which has a counterpart in the U4atac-U6atac snRNA duplex. Biochimie 79: 387–395. doi:10.1016/s0300-9084(97)86148-2

Jukes TH, Cantor CR. 1969. CHAPTER 24 - Evolution of Protein Molecules. In Mammalian Protein Metabolism (ed. H. Munro), Vol. III of, pp. 21–132.

Kandels-Lewis S, Séraphin B. 1993. Involvement of U6 snRNA in 5’ splice site selection. Science 262: 2035–9. doi:10.1126/science.8266100

Kim DH, Rossi JJ. 1999. The first ATPase domain of the yeast 246-kDa protein is required for in vivo unwinding of the U4/U6 duplex. RNA 5: 959–71. doi:10.1017/s135583829999012x

Kuhn AN, Brow DA. 2000. Suppressors of a cold-sensitive mutation in yeast U4 RNA define five domains in the splicing factor Prp8 that influence spliceosome activation. Genetics 155: 1667–1682. doi:10.1093/genetics/155.4.1667

Kuhn AN, Reichl EM, Brow DA. 2002. Distinct domains of splicing factor Prp8 mediate different aspects of spliceosome activation. Proc Natl Acad Sci U S A 99: 9145–9149. doi:10.1073/pnas.102304299

Laggerbauer B, Achsel T, Lührmann R. 1998. The human U5-200kD DEXH-box protein unwinds U4/U6 RNA duplices in vitro. Proc Natl Acad Sci U S A 95: 4188–92. doi:10.1073/pnas.95.8.4188

Lesser CF, Guthrie C. 1993. Mutations in U6 snRNA that alter splice site specificity: implications for the active site. Science 262: 1982–1988. doi:10.1126/science.8266093

Letunic I, Bork P. 2024. Interactive Tree of Life (iTOL) v6: Recent updates to the phylogenetic tree display and annotation tool. Nucleic Acids Res 52: W78–W82. doi:10.1093/nar/gkae268

Madeira F, Madhusoodanan N, Lee J, Eusebi A, Niewielska A, Tivey ARN, Lopez R, Butcher S. 2024. The EMBL-EBI Job Dispatcher sequence analysis tools framework in 2024. Nucleic Acids Res 52: W521– W525. doi:10.1093/nar/gkae241

Madhani HD, Bordonné R, Guthrie C. 1990. Multiple roles for U6 snRNA in the splicing pathway. Genes Dev 4: 2264–2277. doi:10.1101/gad.4.12b.2264

Madhani HD, Guthrie C. 1992. A novel base-pairing interaction between U2 and U6 snRNAs suggests a mechanism for the catalytic activation of the spliceosome. Cell 71: 803–817. doi:10.1016/0092-8674(92)90556-r

Mefford MA, Staley JP. 2009. Evidence that U2/U6 helix I promotes both catalytic steps of pre-mRNA splicing and rearranges in between these steps. RNA 15: 1386–97. doi:10.1261/rna.1582609

Mozaffari-Jovin S, Santos KF, Hsiao HH, Will CL, Urlaub H, Wahl MC, Lührmann R. 2012. The Prp8 RNase H-like domain inhibits Brr2-mediated U4/U6 snRNA unwinding by blocking Brr2 loading onto the U4 snRNA. Genes Dev 26: 2422–2434. doi:10.1101/gad.200949.112

Myslinski E, Branlant C. 1991. A phylogenetic study of U4 snRNA reveals the existence of an evolutionarily conserved secondary structure corresponding to “free” U4 snRNA. Biochimie 73: 17–28. doi:10.1016/0300-9084(91)90069-d

Nguyen THD, Galej WP, Bai X-C, Oubridge C, Newman AJ, Scheres SHW, Nagai K. 2016. Cryo-EM structure of the yeast U4/U6.U5 tri-snRNP at 3.7 Å resolution. Nature 530: 298–302. doi:10.1038/nature16940

Nielsen KH, Staley JP. 2012. Spliceosome activation: U4 is the path, stem I is the goal, and Prp8 is the keeper. Let’s cheer for the ATPase Brr2! Genes Dev 26: 2461–2467. doi:10.1101/gad.207514.112

Noble SM, Guthrie C. 1996. Identification of novel genes required for yeast pre-mRNA splicing by means of cold-sensitive mutations. Genetics 143: 67–80. doi:10.1093/genetics/143.1.67

Ontiveros-Palacios N, Cooke E, Nawrocki EP, Triebel S, Marz M, Rivas E, Griffiths-Jones S, Petrov AI, Bateman A, Sweeney B. 2024. Rfam 15: RNA families database in 2025. Nucleic Acids Res 1–10. doi:10.1093/nar/gkae1023

Plaschka C, Lin PC, Nagai K. 2017. Structure of a pre-catalytic spliceosome. Nature 546: 617–621. doi:10.1038/nature22799

Raghunathan PL. 1998. A Spliceosomal Recycling Factor That Reanneals U4 and U6 Small Nuclear Ribonucleoprotein Particles. Science 279: 857–860. doi:10.1126/science.279.5352.857

Raghunathan PL, Guthrie C. 1998. RNA unwinding in U4/U6 snRNPs requires ATP hydrolysis and the DEIH-box splicing factor Brr2. Curr Biol 8: 847–55. doi:10.1016/s0960-9822(07)00345-4

Reuter JS, Mathews DH. 2010. RNAstructure: Software for RNA secondary structure prediction and analysis. BMC Bioinformatics 11: 1–9. doi:10.1186/1471-2105-11-129

Rodgers ML, Didychuk AL, Butcher SE, Brow DA, Hoskins AA. 2016. A multi-step model for facilitated unwinding of the yeast U4/U6 RNA duplex. Nucleic Acids Res 44: 10912–10928. doi:10.1093/nar/gkw686

Semlow DR, Blanco MR, Walter NG, Staley JP. 2016. Spliceosomal DEAH-Box ATPases Remodel Pre-mRNA to Activate Alternative Splice Sites. Cell 164: 985–998. doi:10.1016/j.cell.2016.01.025

Shannon KW, Guthrie C. 1991. Suppressors of a U4 snRNA mutation define a novel U6 snRNP protein with RNA-binding motifs. Genes Dev 5: 773–785. doi:10.1101/gad.5.5.773

Shuster EO, Guthrie C. 1990. Human U2 snRNA can function in pre-mRNA splicing in yeast. Nature 345: 270–273. doi:10.1038/345270a0

Sikorski RS, Hieter P. 1989. A system of shuttle vectors and yeast host strains designed for efficient manipulation of DNA in Saccharomyces cerevisiae. Genetics 122: 19–27. doi:10.1093/genetics/122.1.19

Small EC, Leggett SR, Winans AA, Staley JP. 2006. The EF-G-like GTPase Snu114p Regulates Spliceosome Dynamics Mediated by Brr2p, a DExD/H Box ATPase. Mol Cell 23: 389–399. doi:10.1016/j.molcel.2006.05.043

Staley JP, Guthrie C. 1999. An RNA switch at the 5’ splice site requires ATP and the DEAD box protein Prp28p. Mol Cell 3: 55–64. doi:10.1016/s1097-2765(00)80174-4

Stevens SW, Abelson J. 2002. Yeast pre-mRNA splicing: Methods, mechanisms, and machinery. Methods Enzymol 351: 200–220. doi:10.1016/s0076-6879(02)51849-8

Stotz A, Linder P. 1990. The ADE2 gene from Saccharomyces cerevisiae: sequence and new vectors. Gene 95: 91–98. doi:10.1016/0378-1119(90)90418-q

Sun JS, Manley JL. 1995. A novel U2-U6 snRNA structure is necessary for mammalian mRNA splicing. Genes Dev 9: 843–854. doi:10.1101/gad.9.7.843

Tagwerker C, Zhang H, Wang X, Larsen LSZ, Lathrop RH, Hatfield GW, Auer B, Huang L, Kaiser P. 2006. HB tag modules for PCR-based gene tagging and tandem affinity purification in Saccharomyces cerevisiae. Yeast 23: 623–632. doi:10.1002/yea.1380

Theuser M, Höbartner C, Wahl MC, Santos KF. 2016. Substrate-assisted mechanism of RNP disruption by the spliceosomal Brr2 RNA helicase. Proc Natl Acad Sci U S A 113: 7798–803. doi:10.1073/pnas.1524616113

Toor N, Keating K, Taylor S, Pyle A. 2008. Crystal structure of a self-spliced group II intron. Science 320: 77. doi:10.1126/science.1153803

Townsend C, Leelaram MN, Agafonov DE, Dybkov O, Will CL, Bertram K, Urlaub H, Kastner B, Stark H, Luhrmann R. 2020. Mechanism of protein-guided folding of the active site U2 / U6 RNA during spliceosome activation. Science 3753: 1–23. doi:10.1126/science.abc3753

Wan R, Bai R, Yan C, Lei J, Shi Y. 2019. Structures of the Catalytically Activated Yeast Spliceosome Reveal the Mechanism of Branching. Cell 177: 339–351.e13. doi:10.1016/j.cell.2019.02.006

Wilkinson ME, Charenton C, Nagai K. 2020. RNA Splicing by the Spliceosome. Annu Rev Biochem 359–388. doi:10.1146/annurev-biochem-091719-064225

Will CL, Lührmann R. 2011. Spliceosome structure and function. Cold Spring Harb Perspect Biol 3: 1–25. doi:10.1101/cshperspect.a003707

Xu D, Nouraini S, Field D, Tang SJ, Friesen JD. 1996. An RNA-dependent ATPase associated with U2/U6 snRNAs in pre-mRNA splicing. Nature 381: 709–13. doi:10.1038/381709a0

Yan C, Wan R, Bai R, Huang G, Shi Y. 2016. Structure of a yeast activated spliceosome at 3.5 Å resolution. Science 353: 904–911. doi:10.1126/science.aag0291

Zhan X, Yan C, Zhang X, Lei J, Shi Y. 2018. Structures of the Human Pre-catalytic Spliceosome and Its Precursor Spliceosome. Cell Res 1–12. doi:10.1038/s41422-018-0094-7

Zhang Z, Kumar V, Dybkov O, Will CL, Zhong J, Ludwig SEJ, Urlaub H, Kastner B, Stark H, Lührmann R. 2024. Structural insights into the cross-exon to cross-intron spliceosome switch. Nature 630: 1012– 1019. doi:10.1038/s41586-024-07458-1

Zhao C, Bellur DL, Lu S, Zhao F, Grassi MA, Bowne SJ, Sullivan LS, Daiger SP, Chen LJ, Pang CP, et al. 2009. Autosomal-Dominant Retinitis Pigmentosa Caused by a Mutation in SNRNP200, a Gene Required for Unwinding of U4/U6 snRNAs. Am J Hum Genet 85: 617–627. doi:10.1016/j.ajhg.2009.09.020

Zhong Z, Yan M, Sun W, Wu Z, Han L, Zhou Z, Zheng F, Chen J. 2016. Two novel mutations in PRPF3 causing autosomal dominant retinitis pigmentosa. Sci Rep 6: 1–6. doi:10.1038/srep37840

