## Supplementary material for "Brr2p-mediated unwinding of U4/U6 is promoted by a mutually exclusive intra-molecular stem loop in U4 and involves destabilization of the 5’ stem-loop of U4": Nielsen-Suppl-all

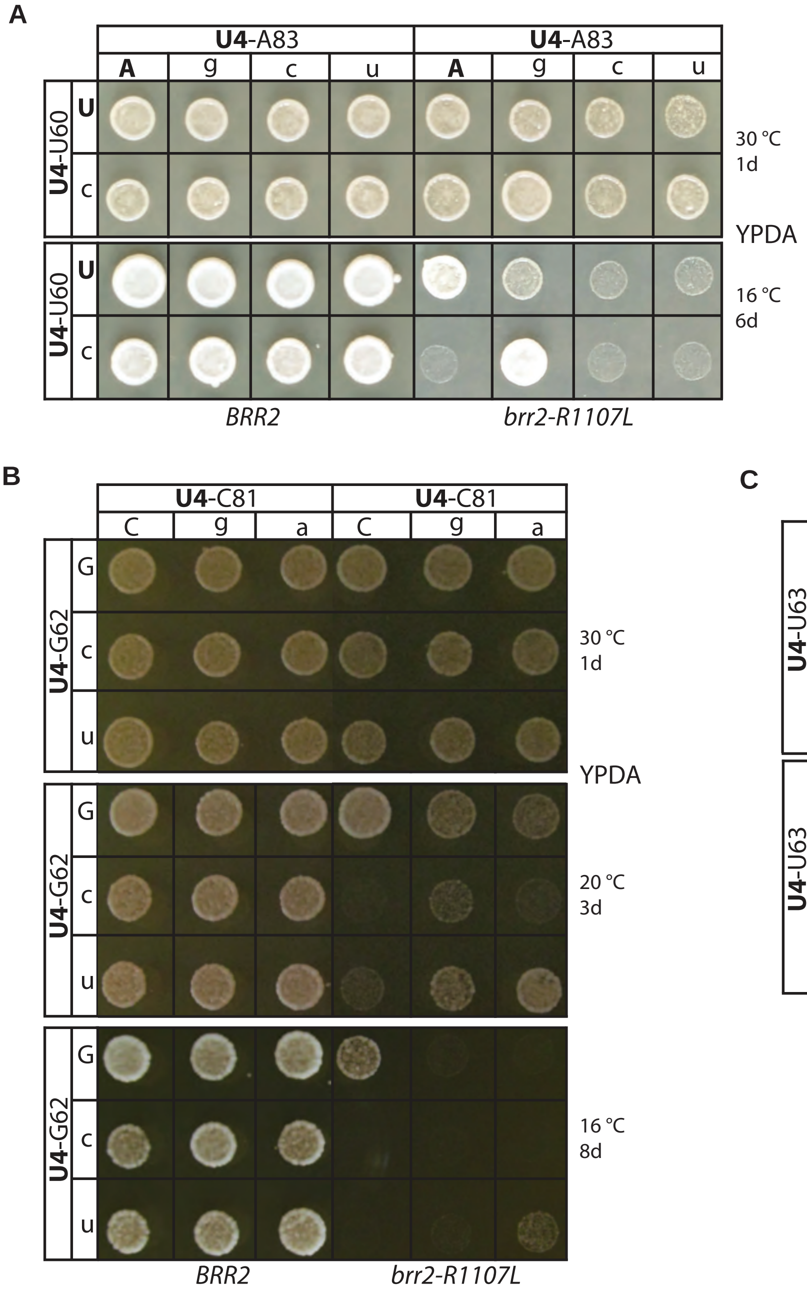


**Supplemental Figure 1.** A novel intra-molecular U4 stem loop, mutually exclusive with U4/U6 stem I, cooperates with *BRR2*. (A-C) Compensatory analysis of U4-ISL1 base pairs U60/A83 (A), G62/C81 (B), and U63/A80 (C), each providing further evidence that U4-ISL1 cooperates with *BRR2*. Point mutations in either base of each base pair exacerbates the growth of *brr2-R1107L*; compensatory combinations that restored base pairing of U4-ISL1 restore growth of *brr2-R1107L*. Growth was assessed and displayed as in Figure 1C on YPDA media.

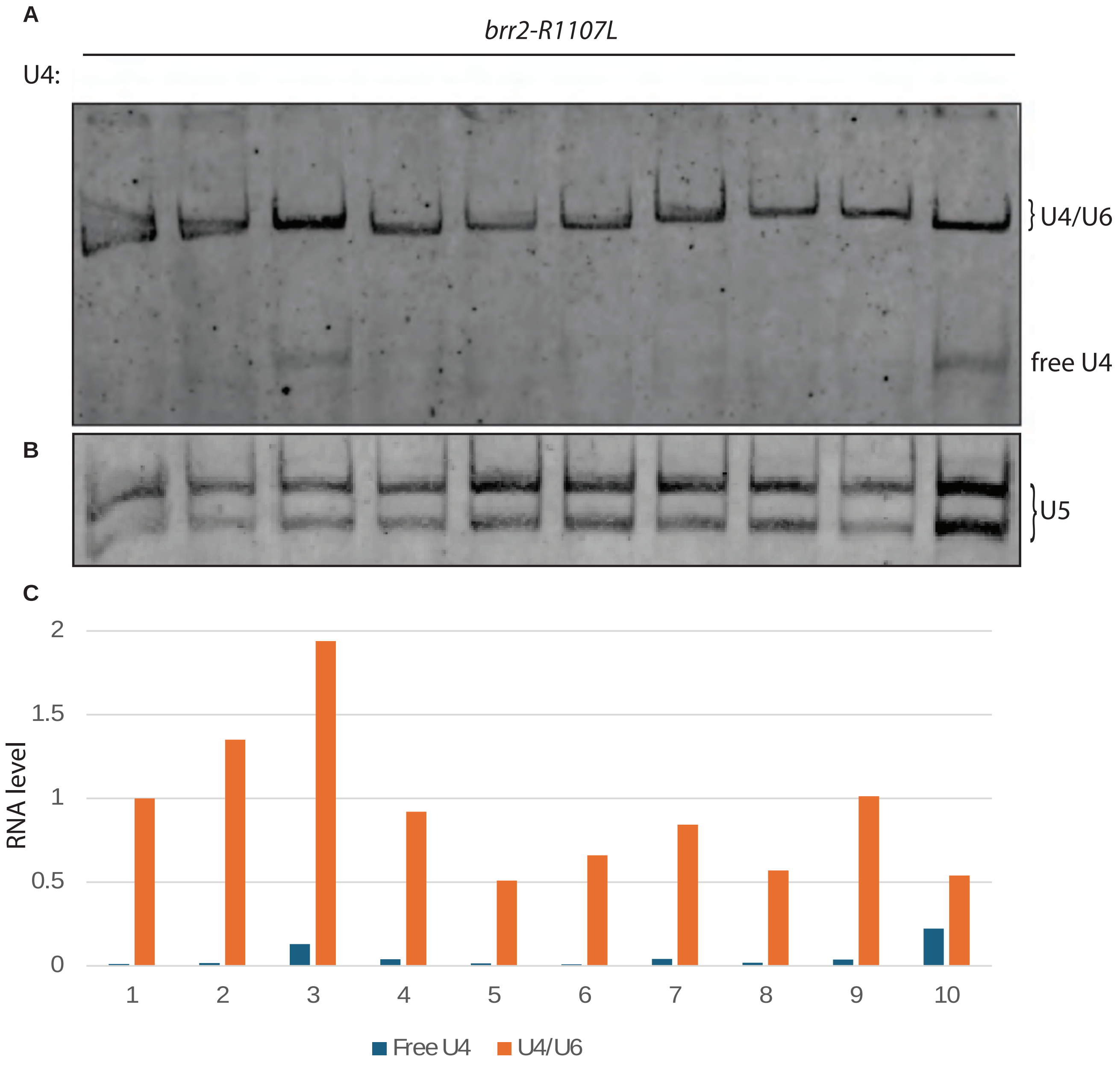


**Supplemental Figure 2.** Mutations in U4 within U4/U6 stem I do not substantially impact U4/U6 steady-state levels, indicating that these mutations are functional for U4/U6 annealing. (A) The *brr2-R1107L* mutant strain with the indicated U4 mutations were grown in rich YPDA media at 30 °C to a $OD_{600}$ of 0.33, after which the culture was shifted to 16 °C for 2 hrs. Cold phenol extracted RNA was separated on a native 8% PAGE gel in 1x TBE buffer and transferred to a membrane and probed by northern blotting against U4 snRNA. (B) As in (A) but probed for U5 snRNA. (C) Quantitation of bands in panels A and B to determine the fraction of free U4 snRNA levels, relative to total U4 snRNA levels (the sum of free U4 snRNA and base paired U4/U6) and the total U4/U6 levels, relative to U5 snRNA levels; for both calculations, all values were normalized to the levels in the strain with WT U4 snRNA. Aside from G58C, which confers a cold-sensitive phenotype in a WT *BRR2* strain, only G58A showed a clear, free U4 snRNA band, but the levels of U4/U6 snRNA nevertheless remained high in this strain. The U4/U6 levels for all mutations were within 2-fold of wild-type U4/U6 levels; there is a slight trend of the mutations that suppress *brr2-R1107L* (lanes 2-4) showing increased U4/U6 snRNA levels and of the mutations that exacerbate *brr2-R1107L* (lanes 5-8) showing reduced U4/U6 snRNAs consistent with the reduce tri-snRNPs observed for the 61 mutant in Figure 5B. However, disrupting U4-ISL1 impacts Brr2p-mediated unwinding of U4/U6 independent of any impact on U4/U6 or tri-snRNP levels (Figure 5). It is not clear why U4-G58C has a more pronounced growth defect in WT *BRR2*, but it may be due to an annealing defect; the mutation has previously been shown to accumulate a splicing complex prior to pre-B complex formation (Hu et al. 1995), consistent with a defect in regenerating tri-snRNPs.

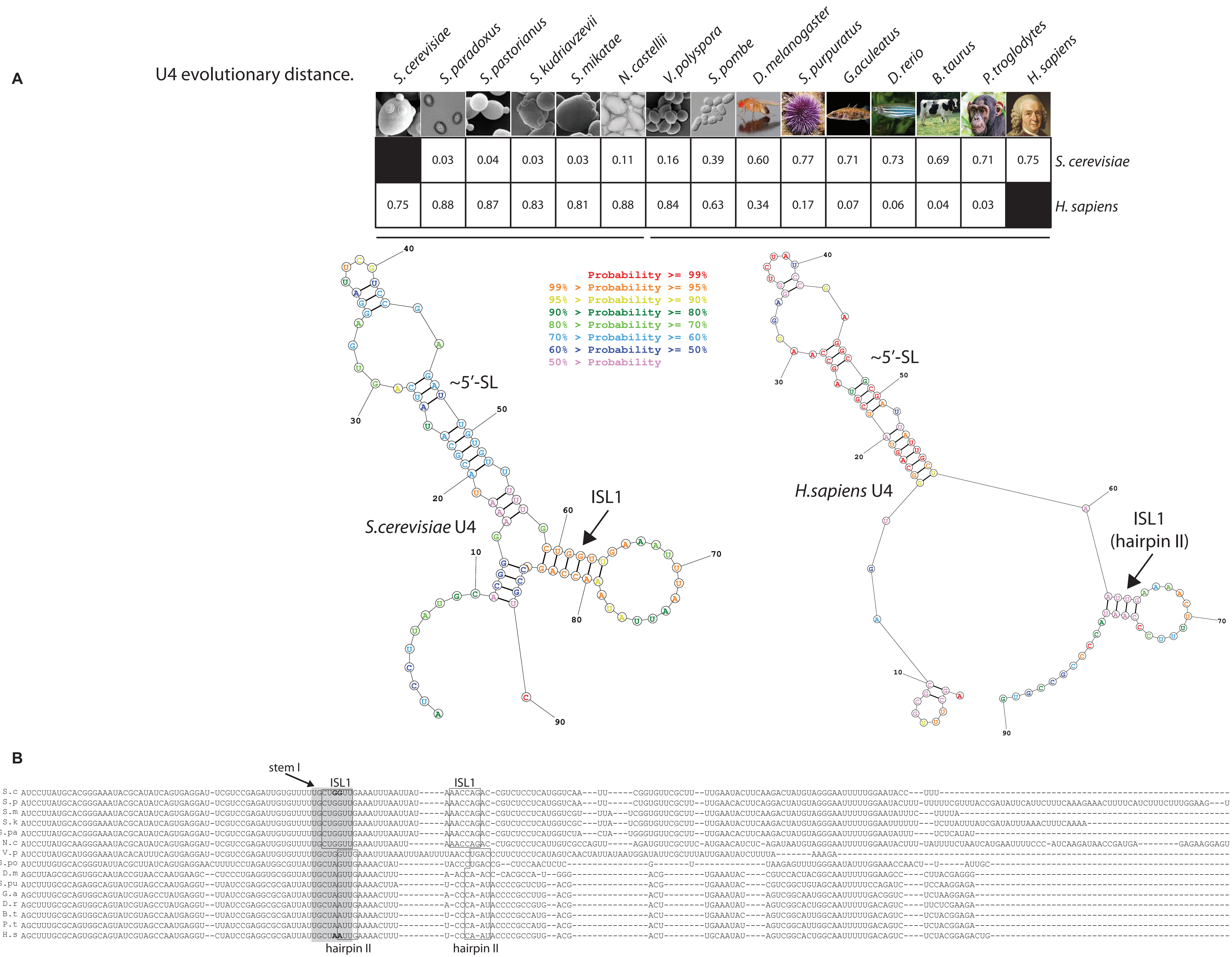
A
U4 evolutionary distance.
S.cerevisiae
S.paradoxus
S.pastorianus
S.kudriavzevii
S.mikatae
N.castellii
V.polyspora
S.pombe
D.melanogaster
S.purpuratus
G.aculeatus
D.rerio
B.taurus
P.troglodytes
H.sapiens
S. cerevisiae: 0.03 0.04 0.03 0.03 0.11 0.16 0.39 0.60 0.77 0.71 0.73 0.69 0.71 0.75
H. sapiens: 0.75 0.88 0.87 0.83 0.81 0.88 0.84 0.63 0.34 0.17 0.07 0.06 0.04 0.03
Probability >= 99%
99% > Probability >= 95%
95% > Probability >= 90%
90% > Probability >= 80%
80% > Probability >= 70%
70% > Probability >= 60%
60% > Probability >= 50%
50% > Probability
~5′-SL
S.cerevisiae U4
ISL1
H.sapiens U4
ISL1 (hairpin II)
B
stem I
ISL1
hairpin II
S.c
S.p
S.m
S.k
S.pa
N.c
V.p
S.po
D.m
S.pu
G.a
D.r
B.t
P.t
H.s

**Supplemental Figure 3.** Evidence that U4-ISL1 formation and function in preventing reannealing of stem I is conserved across eukaryotes. (A) The evolutionary distance of U4 between either *H. sapiens* or *S. cerevisiase* and the indicated species is shown. Evolutionary distance was calculated by the Jukes-Cantor model (Jukes and Cantor 1969). The predicted structure (Reuter and Mathews 2010) of U4 nts. 1-90 from *S. cerevisiase* (left) and *H. sapiens* (right) with either U4-ISL1 or the related hairpin II indicated, respectively. An elongated 5'-SL, just two nucleotides upstream of the 5'SL, is predicted as well. The nucleotides are color coded according to the key indicating the likelihood of the proposed base-pairs. (B) Clustal Omega alignment of the U4 seed sequences from Rfam showing the species used in panel A. The U4/U6 stem I sequence is colored grey. The two GG nucleotides in S. c. that correspond to two AA nucleotides in H. s. in stem I are in bold. The top-listed six species can all form U4-ISL1 (underlined in panel A on the left, below the evolutionary distance). The nucleotides that form the left side of U4-ISL1 are boxed and fully overlapping with U4/U6 stem I, and the downstream sequence that forms the right side of ISL1 is boxed. The species below N. c. all have the potential to form hairpin II (underlined in panel A on the right below the evolutionary distance). The nucleotides that form the left side of hairpin II are boxed and the downstream sequence that forms the right side of hairpin II is boxed. As with ISL1, hairpin II overlaps with and is consequently mutually exclusive with U4/U6 stem I, consistent with a role for hairpin II, as for ISL1, in preventing U4/U6 stem I reannealing, after Brr2p-mediated unwinding.

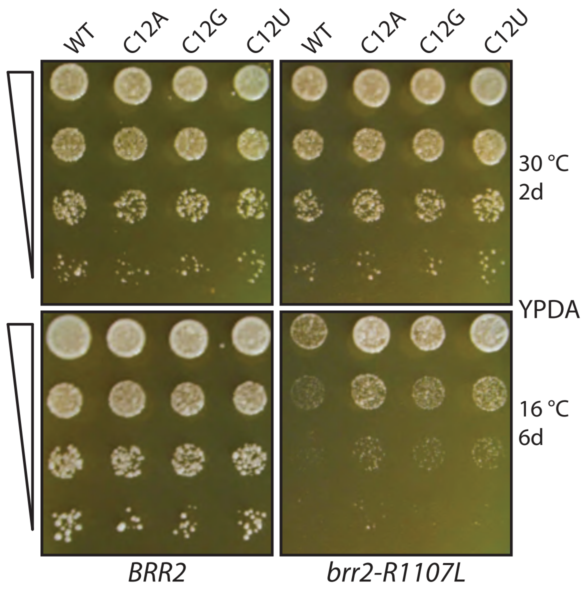


**Supplemental Figure 4.** Non-allelic-specific suppression of brr2-R1107L by mutations at position U4-C12. Growth was assessed and displayed as in Figure 1B.

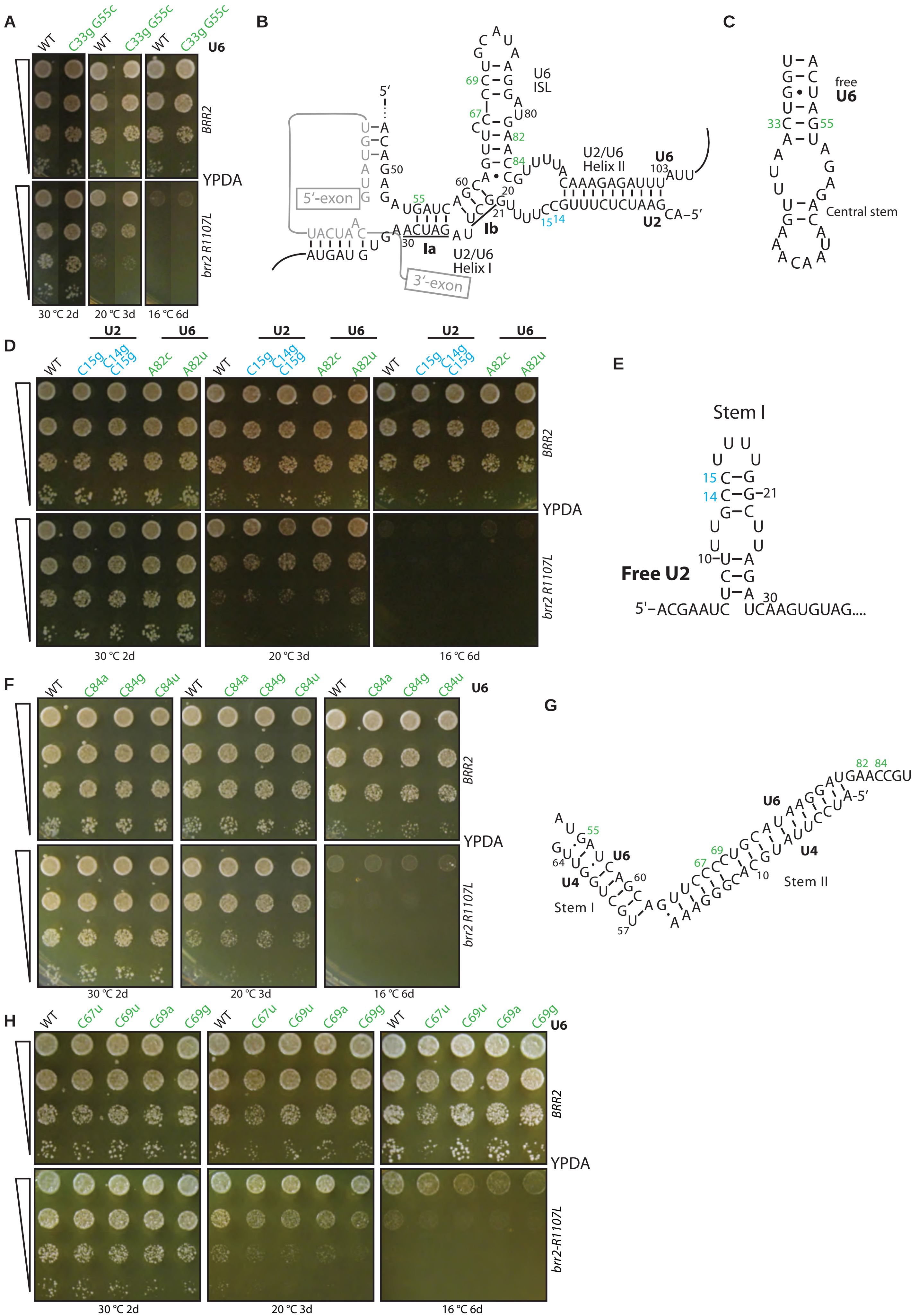
A
WT
C33g G55c
U6
BRR2
YPDA
brr2 R1107L
30 °C 2d
20 °C 3d
16 °C 6d
B
5′
5′-exon
3′-exon
U6 ISL
U2/U6 Helix II
U2/U6 Helix I
Ia
Ib
U6
U2
CA–5′
C
free U6
Central stem
D
U2
U6
WT
C15g
C14g C15g
A82c
A82u
BRR2
YPDA
brr2 R1107L
30 °C 2d
20 °C 3d
16 °C 6d
E
Stem I
Free U2
5′–ACGAAUC UCAAGUGUAG....
F
WT
C84a
C84g
C84u
U6
BRR2
YPDA
brr2 R1107L
30 °C 2d
20 °C 3d
16 °C 6d
G
U6
U4
Stem I
Stem II
A-5′
H
WT
C67u
C69u
C69a
C69g
U6
BRR2
YPDA
brr2-R1107L
30 °C 2d
20 °C 3d
16 °C 6d

**Supplemental Figure 5.** Mutations that alter the stability of U2/U6 helix Ia, U2 stem I, the U6 central stem, or the U6-ISL do not interact genetically with the *brr2-R1107L* mutation. (A) Tests for genetic interactions between U2/U6 helix Ia and *brr2-R1107L*. The mutation U6-U55C, which destabilizes U2/U6 helix Ia, failed to exacerbate *brr2-R1107L*. A second site, compensatory mutation (C33G) in U6 was included to maintain base pairing in the mutually exclusive U6 central stem (Fortner et al. 1994), illustrated in panel C. The data from each temperature is from the same plate, but the image was cut and spliced to juxtapose the relevant mutants. (B) Schematic representation of U2/U6 with pre-mRNA bound and with residues that were mutated and analyzed in this figure; these residues are numbered and colored blue (U2) or green (U6). (C) Schematic representation of the central stem in free U6 with mutated residues number and colored green. (D) Tests for genetic interactions between U2 stem loop I or the U6-ISL and *brr2-R1107L*. U2-C14G and U2-C14G/C15G both destabilize U2 stem I and thereby favor formation of U2/U6 helix I. If formation of U2/U6 helix I, like U4 ISL1, promotes U4/U6 unwinding, then these mutations should suppress *brr2-R1107L*. Neither mutation genetically interacts with *brr2-R1107L*. The U6 G82C and G82U mutations destabilize the U6-ISL without impacting U4/U6 stem II. If the U6-ISL, like U4-ISL1, promotes U4/U6 unwinding, then these mutations should exacerbate brr*2-R1107L*. Neither mutation exacerbates *brr2-R1107L*. (E) Schematic representation of the structure of stem I of U2 with mutated residues numbered and colored blue. (F) More tests for genetic interactions between the U6 ISL1 and *brr2-R1107L*. U6-C84 mutations, which destabilize the U6-ISL without impacting U4/U6 stem II, do not exacerbate *brr2-R1107L*, as we would have expected if the U6 ISL, like U4-ISL1, promoted U4/U6 unwinding. (G) Schematic representation of U4/U6 stem I and stem II with mutated residues numbered and colored green. (H) Yet more tests for genetic interactions between the U6 ISL1 and *brr2-R1107L*. Mutations at U6-C67 disrupt the U6 ISL as well as U4/U6 stem II, whereas mutations at U6-C69, at the bulge in U4/U6 stem II, disrupt the U6 ISL but do not disrupt U4/U6 stem II, although C69G might hyperstabilize U4/U6 stem II. No mutation at either position exacerbated *brr2-R1107L*, as we would have expected if the U6 ISL, like U4-ISL1, promoted U4/U6 unwinding. The negative results with the two C67 mutations might be explained by offsetting effects resulting from destabilization of both the U6 ISL and U4/U6 stem II, but all of the other eight mutations tested, which specifically destabilized the U6 ISL, did not genetically interact with *brr2-R1107L*. In wild type *BRR2*, U6-C67u exhibited a weak cs phenotype at lower temperatures; this phenotype may be due to a U4/U6 annealing defect, because it is base paired with U4-G14, which when mutated also displays a strong cs phenotype that is correlated with an annealing defect (Shannon and Guthrie 1991). In all panels, growth was assessed by frogging of a dilution series of the indicated yeast strains. Growth was assessed as in Figure 1B.

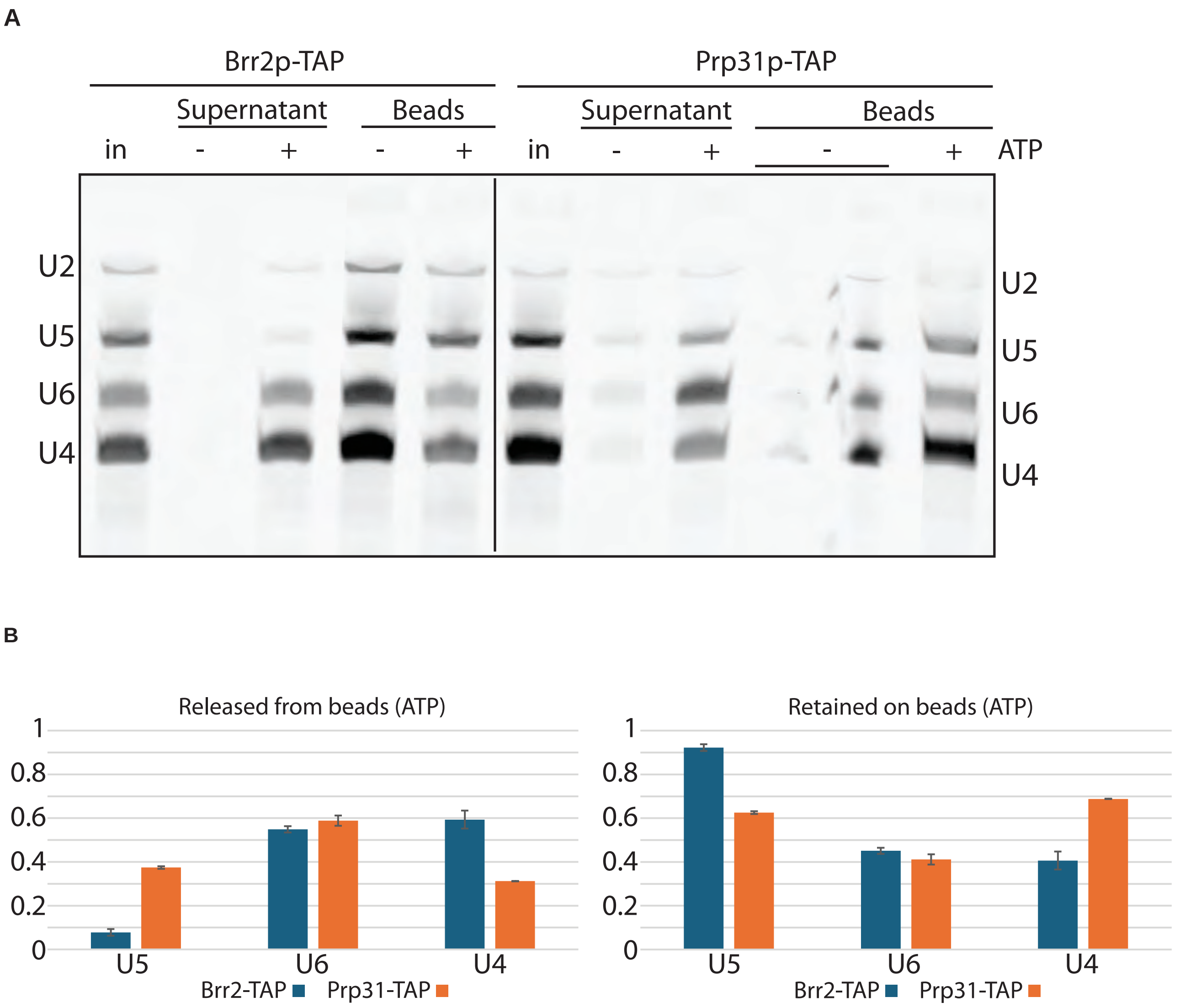


**Supplemental Figure 6.** Evidence that Prp31p readily rebinds U4 snRNA after release by Brr2p. (A) Dissociation of snRNAs from Brr2p or Prp31p were executed as described in Figure 6D but omitting the U4 5′SL oligo. (B) Quantitation of panel A showing the fraction of U4, U5, or U6, after incubation of Brr2p-TAP or Prp31p-TAP tri-snRNPs in the presence of ATP, that was released from (left) or retained bound (right) to the IgG beads. SD is shown and n=3 as in Figure 6E. In the absence of a U4-5′SL RNA competitor but in the presence of ATP, the same amount of U6 is observed in the supernatant for Prp31p-TAP tri-snRNPs and Brr2p-TAP tri-snRNPs, but only half as much U4 is observed in the supernatant for Prp31p-TAP tri-snRNPs as compared to Brr2p-TAP tri-snRNPs. By contrast, in the presence of a U4-5′SL RNA competitor and ATP, the same amount of U6 and U4 is observed in the supernatant for Prp31p-TAP tri-snRNPs and Brr2p-TAP tri-snRNPs (Figure 6E). Consequently, we infer that in the experiment shown in this figure, Prp31p dissociates from both U6 and U4 upon Brr2p-mediated U4/U6 unwinding, but in the absence of competitor, Prp31p rebinds U4. The reaction containing Prp31p-TAP in the absence of ATP was mistakenly loaded across two lanes as indicated.
